# Modular engineering of an extracellular contractile injection system for protein delivery to mammalian and bacterial cells

**DOI:** 10.64898/2026.05.27.728243

**Authors:** Charles F. Ericson, David Schaumann, Elena R. Toenshoff, Pavel Afanasyev, Martin Pilhofer

## Abstract

Extracellular contractile injection systems (eCIS) are autonomous, proteinaceous macromolecular machines, capable of injecting diverse payloads into target cells and thus, promising for biomedical applications. Central mechanistic aspects such as cargo loading are poorly understood and only a few examples of functional modifications have been reported.

Here, we present “<u>pro</u>grammable <u>CIS</u> (PROCIS),” a versatile delivery platform based on engineered eCIS from *Algoriphagus machipongonensis*. Through structure-guided engineering of the tail fiber, we redirect PROCIS toward mammalian or bacterial targets. Furthermore, we identify a conserved AAA+ family ATPase as a required factor for cargo loading into the inner tube lumen. We characterize an N-terminal cargo loading sequence, which is critical for the loading of cargo and can be repurposed for directing non-native cargos to the tube lumen. Additionally, we demonstrate that a tape measure protein acts as a molecular ruler, allowing for precise tuning of particle length that correlates with cargo capacity. Finally, we integrate our insights and demonstrate the functional delivery of bioactive proteins such as Cre recombinase and beta-lactamase to *E. coli* and HeLa cells, respectively.

Our results provide a framework for understanding the eCIS mechanism and establish PROCIS as a programmable toolkit for the targeted intracellular delivery of large biomolecules.

## Introduction

The intracellular delivery of large biomolecules has garnered increasing attention with the rise of protein-based tools for therapeutic and functional use, such as CRISPR-Cas systems, transcription factors, and engineered enzymes. Delivering proteins across the cell membrane remains a major challenge due to their large size and charged nature, which hinder passive membrane (or cell envelope) permeation without a delivery vehicle^1^. Furthermore, achieving precise cell-type specificity with the same protein delivery tool is crucial for maximizing therapeutic efficacy and broadening the range of targetable cell types.

Various strategies have been explored to facilitate protein delivery into cells, including cell-penetrating peptides^2^, synthetic nanocarriers^3^, and viral vectors^4,5^, each characterized by unique mechanisms, efficiencies, and challenges. Recently, bacterially derived protein delivery systems have emerged as promising platforms to address these challenges, leveraging mechanisms for efficient membrane translocation and, in some cases, intrinsic cell-type targeting. Bacterial secretion systems such as the type three secretion system (T3SS)^6^, type four secretion system (T4SS)^7^, and type six secretion system (T6SS)^8^ have been explored for re-engineering as protein delivery platforms, but their dependence on being anchored to the membrane of the producing bacterium necessitates direct cell-to-cell contact of cells for effective cargo transfer, limiting their functionality.

Bacterially produced extracellular contractile injection systems (eCIS), are structurally related to both bacteriophage contractile tails and membrane-bound T6SSs. eCIS function as autonomous, diffusible particles capable of targeting cells from the extracellular space, thereby overcoming the need for direct cell-cell contact. While eCIS, such as the Photorhabdus virulence cassettes (PVCs), have been successfully modified to bind and inject proteins into eukaryotic cells both *in vitro* and *in vivo*, the mechanisms underlying retargeting and cargo loading remain largely uncharacterized^9–11^.

Here we employ AlgoCIS, an eCIS derived from the marine bacterium *Algoriphagus machipongonensis* PR1, as a model system^12^. The availability of high-resolution structures in extended, contracted, as well as multiple intermediate states has allowed for comprehensive understanding of the firing mechanism^13^. These insights, combined with the genetic tractability and the ability to purify the particle directly from its native host, makes AlgoCIS an ideal system for further characterizing eCIS mechanisms and modifications. Here, we re- engineer AlgoCIS as a distinct eCIS-based delivery platform, the “<u>pro</u>grammable <u>CIS</u>, PROCIS,” while advancing our understanding of its modularity and mechanisms.

## Results

### Retargeting of AlgoCIS to bacterial and mammalian cells by engineering the tail fiber

The AlgoCIS operon is 27 kb in total and consists of 18 individual open reading frames, including all the necessary structural and accessory components for proper assembly and function of the system^12^ (Fig.1a/b). To re-engineer the AlgoCIS into a protein delivery tool, we began by modifying its specificity towards selected targets. Given that other closely related CISs also possess flexible tail fibers that confer specificity, such as the T4 bacteriophage long tail fibers and the Pvc13 tail fiber proteins found in PVCs, we chose to begin by modifying the equivalent tail fiber structures found in AlgoCIS^9,14–16^. Our previous structural studies revealed that the organization of tail fibers of AlgoCIS differ significantly compared to other established CISs. Each of the six tail fibers is comprised of a trimer of the tail fiber protein Alg19. Alg19 has four separate domains, which were labeled as the shoulder, neck, middle, and the C-terminal flexible domains (Fig.1c/d). The shoulder, neck, and middle domains wrap around the baseplate cage, forming a larger encompassing structure around the baseplate, which can be observed in tomographic slices of individual free AlgoCIS (Extended Data Fig. 1a, black arrowhead). The C-terminal 531 residues constitute a flexible arm that extends outward from the neck domain (Fig.1 c and Extended Data Fig. 1a, white arrowhead). This observation suggests a notable degree of conformational freedom for the entire C-terminal domain. Moreover, structural and sequence similarity analysis suggests that the shoulder, neck, and middle domains are conserved across related eCIS-like gene clusters such as the thylakoid-anchored tCIS^17^, while the C-terminal region exhibits a higher variability (Extended Data Fig. 1b-d).

**Fig. 1:**
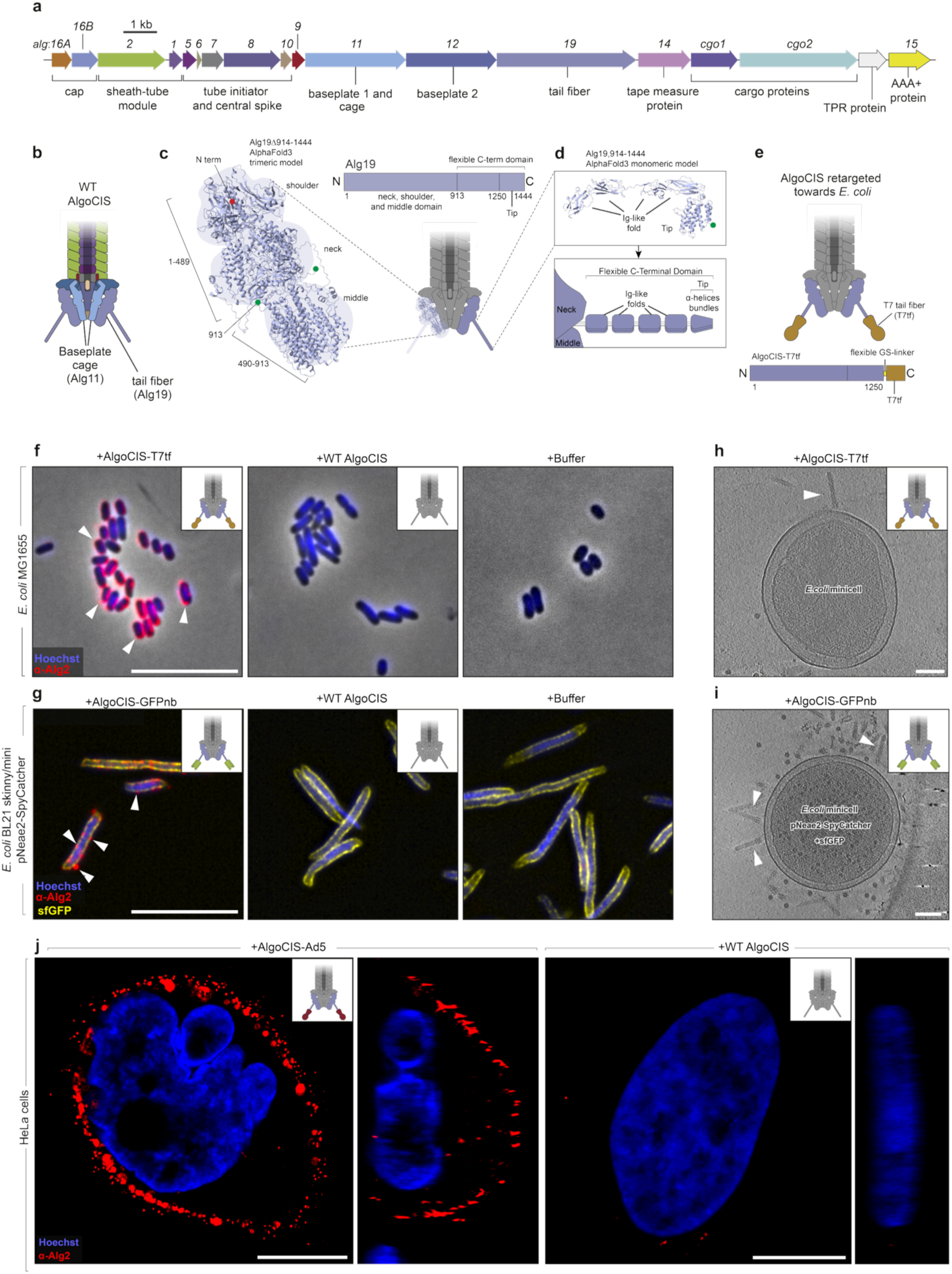
Retargeting of AlgoCIS to bacterial and mammalian cells by engineering the tail fiber. **a.** The AlgoCIS operon from the marine bacterium Algoriphagus machipongonesis PR112 . The operon is 27.3 kb in total and comprised of 18 individual genes that include the structural, cargo, and accessory genes. Functions are shown under the operon, while gene names are shown above. Bar, 1 kb. **b.** Cartoon of WT AlgoCIS baseplate highlighting the orientation and position of the tail fiber protein Alg19. **c.** The AlgoCIS tail fiber protein, Alg19 is 1’444 residues in length and contains four domains, including the neck, shoulder, middle, and a flexible C-terminal domain. A trimeric Alphafold3 model prediction of Alg19 which contains the shoulder, neck and middle domains (Alg19Δ194-1444)^13^, N- and C-termini are shown with red and green dots respectively. **d.** A monomeric Alphafold3 model of the flexible C-terminal domain (residues 914-1’444) of Alg19 reveals four Ig-like folds followed by a bundle of helices, termed as the tip. Below is a schematic representation of the C-terminal flexible domain of Alg19. **e.** Schematic for modifying the Alg19 tail fiber tip by replacing it with the known T7 bacteriophage tail fiber domain, changing the specificity of AlgoCIS towards *E. coli*. **f-g.** Immunofluorescence (IF) light microscopy (LM) images of *E. coli* cells incubated with WT AlgoCIS or AlgoCIS which contains retargeted Alg19 tail fibers (AlgoCIS-T7tf and AlgoCIS-GFPnb). Secondary fluorescent nanobody against the primary antibody against domain IV of the sheath (Alg2) is shown in red. DNA stain Hoechst is shown in blue. sfGFP coating the cells is shown in yellow. AlgoCIS binding to the surface of the *E. coli* is indicated with white arrowheads. Bar, 10 µm.\ **h-i.** Cryo-ET slice of an *E. coli* minicell with retargeted AlgoCIS attached to the surface of the cell. Examples of perpendicularly bound particles are indicated with a white arrowhead. Bar, 100 nm **j.** IF confocal slices of HeLa cells that have been incubated with either WT AlgoCIS or a mutant which retargets the AlgoCIS towards human cells (AlgoCIS-Ad5). Orthogonal views along the Y-Z plane are shown on the right of the images. Bar, 10 µm.

Alg19 has recently been reported being implicated in the rare attachment of the AlgoCIS particles to the surface of the bacterial strain *Echinicola pacifica*, with the flexible C-terminal domain stabilizing and attaching to the bacterial surface^13^. We used an AlphaFold3^18^ prediction of the C-terminal flexible domain, which indicated a structural organization consisting of four Ig-like folds followed by a C-terminal region (residues 1251-1444) forming a bundle of α-helices we termed the “tip” region (Fig.1d). We theorized that by replacing the tail fiber tip with known binding protein domains, we would be able to modify the binding ability of AlgoCIS towards new targets.

We began by replacing the tip of Alg19 with the tail fiber protein from the *E. coli* specific T7 bacteriophage, due to the folding of T7 tail fiber not needing a chaperone^19^ (Fig.1e). The retargeted AlgoCIS were incubated with *E. coli* MG1655 before being analyzed via immunofluorescence (IF) utilizing an anti-Alg2 (sheath) antibody (Fig.1f). We observed strong signal covering the *E. coli* cell surface after co-incubation with the retargeted particles, but not with our WT AlgoCIS control. To expand the re-targeting ability of the system, we also replaced the native C-terminal tip of Alg19 with an anti-GFP nanobody^20^ and incubated the resulting particles (AlgoCIS-GFPnb) with an *E. coli* strain that was coated with sfGFP (Extended Data Fig. 2a/b). IF experiments revealed specific signal on cells incubated with AlgoCIS-GFPnb, while signal was again absent with WT AlgoCIS control (Fig.1g).

To confirm that the observed signals in both assays were indeed due to engineered AlgoCIS particles bound to the cell surface, we set out to image cells by electron microscopy (EM). We utilized an *E. coli* strain (MG1655 Δ*minCDE, mreB-A125V*) that produces *E. coli* minicells with a diameter that allows direct EM imaging without sample thinning^21,22^ (Fig.1h). Negative staining EM and cryo-electron tomography (cryo-ET) imaging of minicells showed a number of engineered AlgoCIS particles bound to the surface of the minicells, while only few particles were seen in the sample that was incubated with WT AlgoCIS (Fig.1h and Extended Data Fig. 2c).

To re-target the system towards human cells, the tail fiber tip of Alg19 was replaced with the adenovirus tip protein which has been shown to naturally bind to the Coxsackievirus and Adenovirus receptor (CAR)^9,23^ (AlgoCIS-Ad5). This strategy for retargeting has previously been shown for PVCs^9^. Upon co-incubation of HeLa cells with AlgoCIS-Ad5, we detected retargeted AlgoCIS on the cell surface via IF, while observing rare unspecific binding events in the sample treated with WT AlgoCIS control (Fig.1j).

In summary, we demonstrate that the C-terminal tip of the Alg19 tail fiber can be replaced with diverse heterologous binding domains, enabling retargeted AlgoCIS particles to bind both bacterial and mammalian targets. Intriguingly, for both approaches, we also screened for a possible cytotoxic effect of engineered AlgoCIS on the target cells and found no significant toxicity (Extended Data Fig. 2d-f). In order to investigate whether target binding also leads to the injection of cargo proteins, we then focused on understanding how cargo is loaded into AlgoCIS particles, followed by loading a non-native cargo that can be detected in the host organism by functional assays.

### Loading sequence mediates loading of native and non-native cargo

Cargo loading in the related eCIS, PVCs, is dependent on an N-terminal loading sequence but the mechanism is unclear^9,24^. We started by analyzing the native AlgoCIS cargo proteins Cgo1 and Cgo2, which have been shown to both load into the inner tube lumen (Fig. 2a)^12^. The analyses with the disorder prediction tool IUpred2a along with AlphaFold3 models indicated that both proteins share an organizational pattern (Fig. 2b/c). Both proteins comprise an N-terminal domain (NTD), followed by a central eCIS effector-associated domain DUF4157, and a C-terminal domain (CTD). Cgo1 and Cgo2 differ significantly in overall size (Cgo1: 54 kDa, 498 residues; Cgo2: 134 kDa, 1’228 residues), arising from length differences in the CTDs. For both cargos, the NTD was predicted to be mostly disordered, in contrast to DUF4157 and CTD (Fig. 2b/c/d). Importantly, other characterized eCIS cargo proteins associated with PVCs (Pdp1 and Pnf1) and MACs (Mif1 and Pne1) revealed that these effectors are lacking DUF4157 but they do exhibit an NTD with a high degree of predicted disorder (Extended Data Fig. 3a). Comparative sequence analyses of the NTDs of the six characterized eCIS cargo proteins showed some sequence similarity between the pairs of cargo from a given system but not between the systems (i.e. similarities between Pdp1 and Pnf1; Mif1 and Pne1; Cgo1 and Cgo2) (Extended Data Fig. 3a-d). In summary, our data indicate that eCIS cargo proteins have a similar architecture with a variable CTD that likely represents the effector domain, an optional DUF4157 domain with a possible function in cargo loading or cargo processing, and a system-specific disordered NTD.

**Fig. 2:**
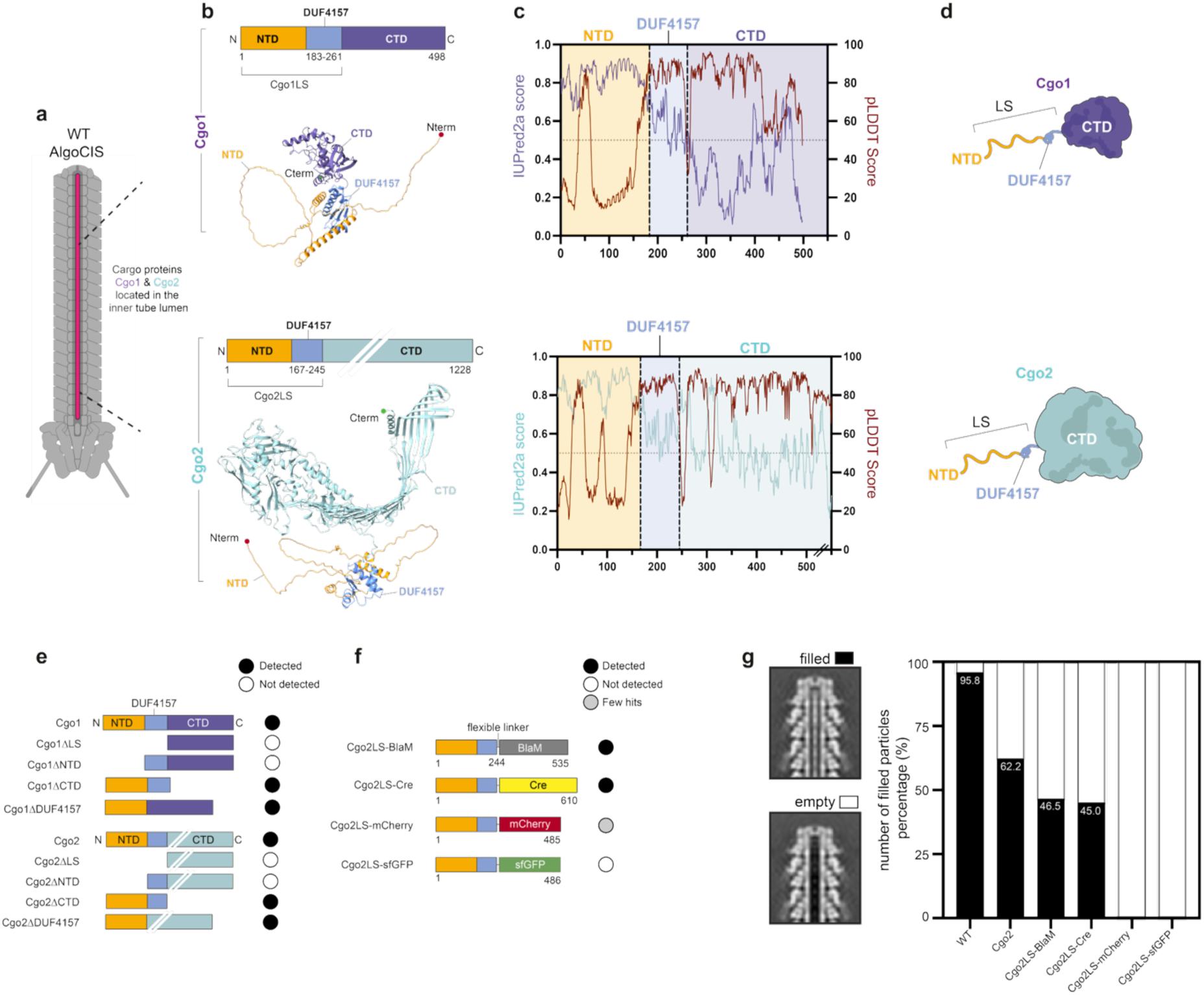
AlgoCIS cargo proteins contain a disordered region that can act as a loading sequence. **a.** Cutaway cartoon of an AlgoCIS particle showing the localization of Cgo1 and Cgo2 within the inner tube lumen. Cargo protein locations are shown in red. **b.** Comparison of Alphafold3 models of both Cgo1 and Cgo2 show a similar domain architecture, which contain a disordered N-terminal domain (NTD) followed by DUF4157, followed by a differing C-terminal domain (CTD). The total length and size of Cgo1 and Cgo2 differ in length and predicted protein size: Cgo1, 498 residues, 54 kDa; Cgo2, 1’228 residues, 133 kDa. **c.** Disorder probability chart from IUPred2A^60^ overlayed with pLDDT confidence score from Alphafold3 structure prediction. Both cargo proteins have a predicted highly disordered NTD followed by DUF4157. The CTD for both proteins varies, but is generally more ordered than the rest of the protein with a higher structural prediction confidence level. **d.** Schematic of the two cargo proteins from AlgoCIS with their conserved N-terminal disordered loading sequences attached to differing C-terminal domains. **e.** Cgo mutant truncation schematics showing deletions of the three domains (NTD, DUF4157, and CTD) in Cgo1 and Cgo2. Samples were purified and sent to MS for determining the cargo loading, filled circles indicate that they were detected in MS of the purified particles while unfilled circles indicate not being detected and grey circles indicate a small number of hits. The results show that the NTDs are critical for cargo loading, with the rest of the Cgo protein not being loaded when the NTD is deleted. **f.** Fusion of distinct reporter proteins to Cgo2LS yielded heterogeneous loading profiles detected through MS. Cgo2LS fused to BlaM or Cre were detected in the purified particles, while sfGFP was not detectable and a few hits were detected for the Cgo2LS-mCherry construct indicating the properties of the reporter might play a factor in cargo loading. **g.** Quantification of cargo-loaded AlgoCIS particles. Purified complexes were imaged by cryo-EM, and the percentage of filled particles was determined through 3D classification by the number of particles corresponding to filled/empty 3D classes (central slices through the 3D classes are presented on the left). This confirms the MS data where the BlaM and Cre were loaded, however the sfGFP and mCherry were not detectable in the tube of the AlgoCIS.

To characterize the regions that are required for cargo loading, we used a *Δcgo1/Δcgo2* double knockout strain to re-introduce individually Cgo1 and Cgo2, respectively, as partial deletion mutants: ΔNTD-DUF4157 (lacking both the NTD and DUF4157, which together hereafter will be referred to as the loading <u>s</u>equence, LS), ΔNTD, ΔDUF4157, and ΔCTD. Mass spectrometry (MS) analysis of purified mutant AlgoCIS particles revealed that neither cargo was detected in both the ΔLS and ΔNTD mutants. In contrast, mutated cargo could be detected by MS for the ΔDUF4157 and ΔCTD mutants. This suggests that the NTD is required for cargo loading, while DUF4157 and CTD are dispensable (Fig. 2e).

Next, we leveraged these insights to load non-native reporters into AlgoCIS particles. We chose Cgo2LS for cloning practicality and fused it to various reporter proteins to test loading of non-native cargos. These included genes encoding sfGFP (26.81 kDa), mCherry (26.71 kDa), β-lactamase (BlaM) (31.52 kDa), and Cre recombinase (38.56 kDa) into a Δ*cgo1*/Δ*cgo2* knockout background. AlgoCIS particles were purified and subsequently analyzed via MS. Excitingly, BlaM and Cre were consistently detected in the respective particle preparations (Extended Data Fig. 4). mCherry was only detected with a very low number of unique peptide counts. sfGFP was not detected in either the Cgo2LS-sfGFP mutant or when fused to the C-terminal end of the native Cgo2 sequence (Fig. 2f and Extended Data Fig. 4).

We assessed the loading efficiency by single-particle cryo-EM images of filled and empty AlgoCIS particles. We found that for the Cgo2LS-BlaM and Cgo2LS-Cre constructs 46.5% and 45.0% of the particles were loaded, respectively (Fig. 2g). This compares to 95.8% of particles in the WT (containing both Cgo1 and Cgo2) and 62.18% of filled particles for single Cgo1 knockout mutants^12^. Interestingly, in the Cgo2LS-mCherry and Cgo2LS-sfGFP constructs we were unable to detect any loaded structures, in line with our MS result. We speculated that the local flexibility of the cargo proteins might be relevant for the distinct cargo loading efficiency. To test this, we estimated Root Mean Square Fluctuation (RMSF) profiles, with BlaM (2.12) and Cre (1.00) showing higher RMSF values than mCherry (0.76) and sfGFP (0.90), indicating greater local flexibility that may correlate with their susceptibility to unfolding. Given that these protein values broadly correlate with the ability to load these proteins into AlgoCIS, suggesting that the ability to load proteins is dependent on their stability and ability to be unfolded.

In summary, we identified that AlgoCIS inner tube cargo loading requires a disordered loading sequence, which can in turn be used to load certain non-native cargos into the system.

### An AAA+ protein mediates cargo loading

Next, we set out to understand the mechanism of cargo loading into the inner tube lumen, which is unknown for any eCIS. Space limitations inside the tube lumen and the large size of known cargo proteins indicate that such cargo must be loaded in an (partially-)unfolded state, which is supported by the fact that the cargo densities cannot be resolved in cryo-EM structures of eCIS particles^25,26^ . Together with our above finding that highly stably folded proteins may not be loaded, this led us to hypothesize that active protein unfolding may be required for loading. We therefore noticed the presence of a gene encoding an AAA+ protein, Alg15, in the AlgoCIS gene cluster (Fig. 1a and Fig. 3a) ^24,27–29^. Notably, many other eCIS operons encode AAA+ homologs; however, their role has thus far not been determined^30–32^. An AlphaFold3 structure prediction of Alg15 revealed a hexameric ring-shaped structure with a central pore, which is an architecture that is typical of AAA+ proteins (Fig. 3b).

**Fig. 3:**
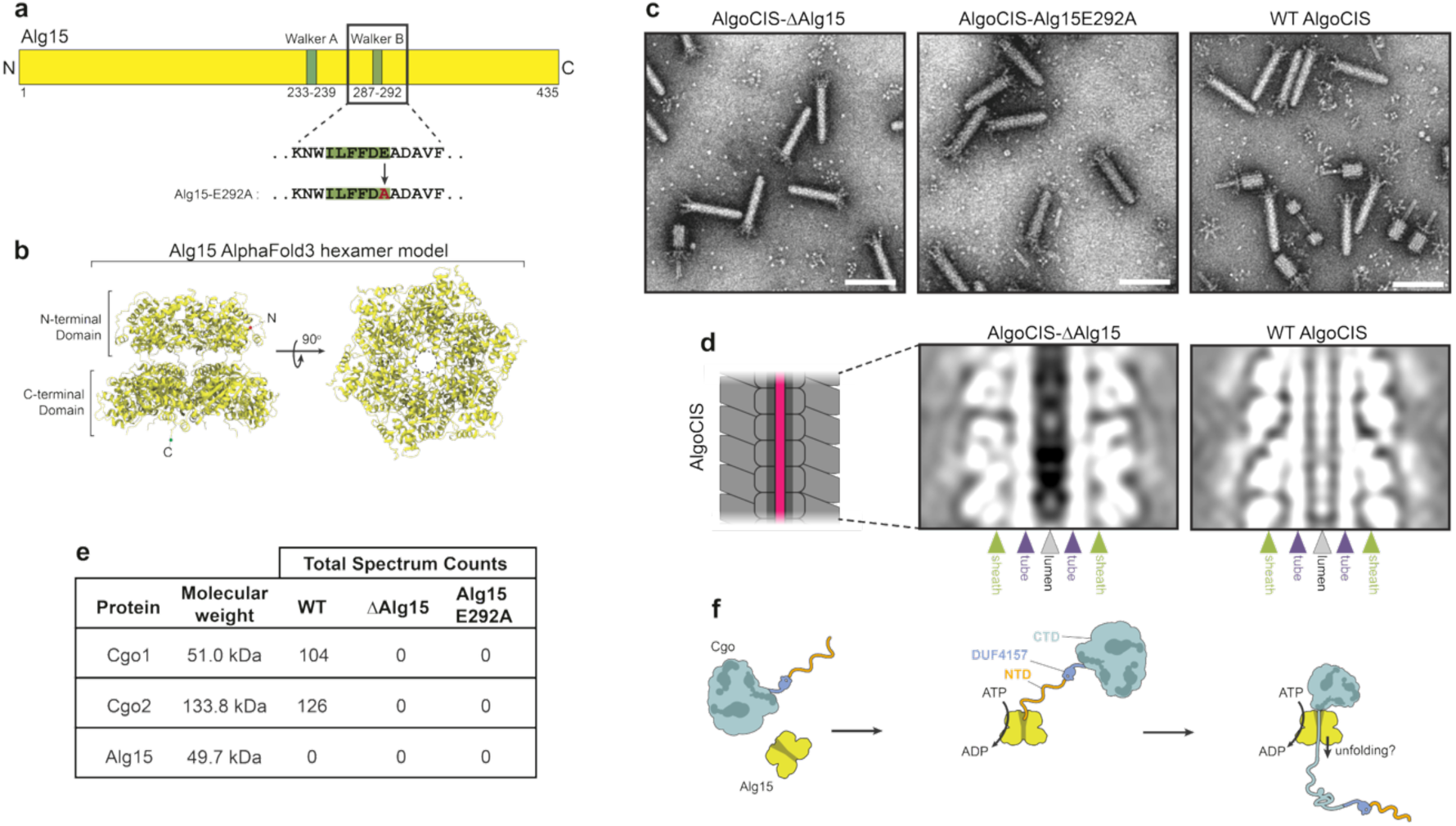
An AAA+ protein mediates cargo loading. **a.** Alg15 contains homology to known AAA+ proteins and contains the Walker A and Walker B motifs that are commonly associated with them. The sequence of the Walker B motif is enlarged to indicate the glutamic acid to alanine point mutation (Alg15 E292A) to stop ATP hydrolysis. **b.** An AlphaFold3 prediction of the hexameric Alg15 shows a stacked ring-like structure typical of other AAA+ proteins. Two separate domains are visible along with a central channel. **c.** Negative stain EM images of purified AlgoCIS from WT, ΔAlg15, and Alg15 A mutants. The ΔAlg15 and Alg15 E292A mutants are still able to produce AlgoCIS particles that appear similar to the WT particles in overall shape and size. Bar, 100 nm. **d.** Cryo-EM single particle analysis of the AlgoCIS sheath-tube modules demonstrate visible densities in the inner tube lumen of the WT AlgoCIS, but a lack of density in the ΔAlg15 particles when the same number of particles from each 3D reconstruction were analyzed. This indicates that while Alg15 is not necessary for assemble of the particle, it is critical for the protein cargo loading. **e.** Chart highlighting MS data from the purified AlgoCIS particles. Neither Cgo1 nor Cgo2 could be detected in the ΔAlg15 or Alg15 E292A particles. Alg15 was also never detected in any of the purified particles (central slice through 3D-reconstruction of distinct 3D classes), indicating that it is not a core structural component and is not associated with the final particle. The lack of Cargo loading in Alg15 E292A indicates that the enzymatic activity of Alg15 is necessary for loading of Cgo1/2. **f.** Suggested cartoon model of ATPase-dependent Alg15 mediated cargo protein unfolding, with Alg15 loading the cargo proteins via the LS of the individual cargo proteins.

To test the role of Alg15 in cargo loading, we generated a deletion (Δ*alg15*) as well as a hydrolysis-deficient mutant (AlgoCIS-Alg15 E292A) by substituting the catalytic glutamic acid with an alanine in the conserved Walker B motif (DExx-motif)^33^. We then purified the AlgoCISs, imaged them using negative staining EM, and found that both mutants still assembled AlgoCIS particles without any obvious major structural differences (Fig. 3c). Further examination of AlgoCIS-ΔAlg15 particles by single particle cryo-EM showed the absence of the luminal density in the inner tube lumen that is seen in WT particles and represents the cargo^12^ (Fig. 3d). Finally, MS confirmed the absence of Cgo1/Cgo2 proteins in purified particles of both mutants (AlgoCIS-ΔAlg15 and AlgoCIS-Alg15 E292A) (Fig. 3e). In agreement with previous cryo-EM structures that did not show Alg15 as an integral structural component of AlgoCIS particles, MS did also not detect any Alg15 hits in any of the analyzed samples, including the WT (Fig. 3e). In conclusion, our data reveal Alg15 as a critical non-structural factor for loading cargo into AlgoCIS particles.

### Tape measure protein controls AlgoCIS length and amount of loaded cargo

With our insights into cargo loading, we sought to determine if changing the AlgoCIS length resulted in altered amounts of cargo per particle. The length determining mechanism utilized by eCIS is not well understood, with some evidence indicating a correlation with the predicted tape measure protein (TMP)^29,34^. In related bacteriophages, the length of TMP is directly correlated with the length of the phage tail^35,36^. This is accomplished by the mostly linear TMP being coated with chaperone proteins and the C-terminal end of the TMP binding to the baseplate during phage assembly. The chaperones are then replaced with the inner tube and sheath proteins, resulting in phage tails of uniform lengths^37^.

By aligning the secondary structure prediction of the AlgoCIS TMP (Alg14) with the TMPs from other eCIS that have known particle lengths, a conserved pattern became evident. Both the N- and C-terminal regions contained a mixture of β-strands and α-helices (Extended Data Fig. 6a). Similar to bacteriophage TMPs, the middle regions of the eCIS TMPs were predicted to almost exclusively be comprised of α-helices but varied in the length and number of α-helices (Extended Data Fig. 5a). When the lengths of known eCIS were plotted against their corresponding TMPs, there was a general correlation between them with an R^2^ of 0.84 (Extended Data Fig. 5b).

Motivated by the correlation between the particle and TMP residue length, we attempted to modify the length of the AlgoCIS particle. By calculating the ratio of residues of Alg14 to the total particle length of WT AlgoCIS (120.6 nm/526 residues), we were able to calculate a coefficient of 0.229 nm per TMP residue^12^ (Fig. 4a/b). Based on the AlphaFold3 predicted structure of Alg14, we selected the resulting highest confidence α-helices found in the middle α-helices-rich region in Alg14 (Extended Data Fig. 5c). This revealed a region of four successive α-helices which contained higher confidence predictive values (residues 163-213). Duplicating these α-helices resulted in an Alg14 with a length of 590 residues (termed Alg14-10%longer), compared to the WT length of 526 residues (Fig. 4c). By multiplying the Alg14-10%longer mutant by our previous calculated coefficient, we estimated that the particles would now be around 132.5 nm, i.e., 10% longer than the ∼121 nm of the WT. Intriguingly, negative staining EM and subsequent 2D classification showed that the AlgoCISAlg14-10%longer mutant exhibited a length of around 138 nm, similar to our predicted value (Fig. 4d). To test the robustness of this strategy, we also created a mutant where we duplicated the entire internal α-helices region of Alg14 (Alg14-30%longer), in addition to an Alg14 mutant where the entire internal α-helices were deleted (Alg14-shorter) (Fig. 4c/d). Based on the coefficient, we predicted that Alg14-30%longer mutant would result in a 152.0 nm long particle, with the Alg14-shorter mutant was predicted to be 85.1 nm in length. The 2D classification analyses of purified mutants showed that the average lengths of the AlgoCIS-Alg14-30%longer and AlgoCIS-Alg14-shorter particles were ∼161 nm and ∼89 nm respectively (Fig. 4d).

**Fig. 4:**
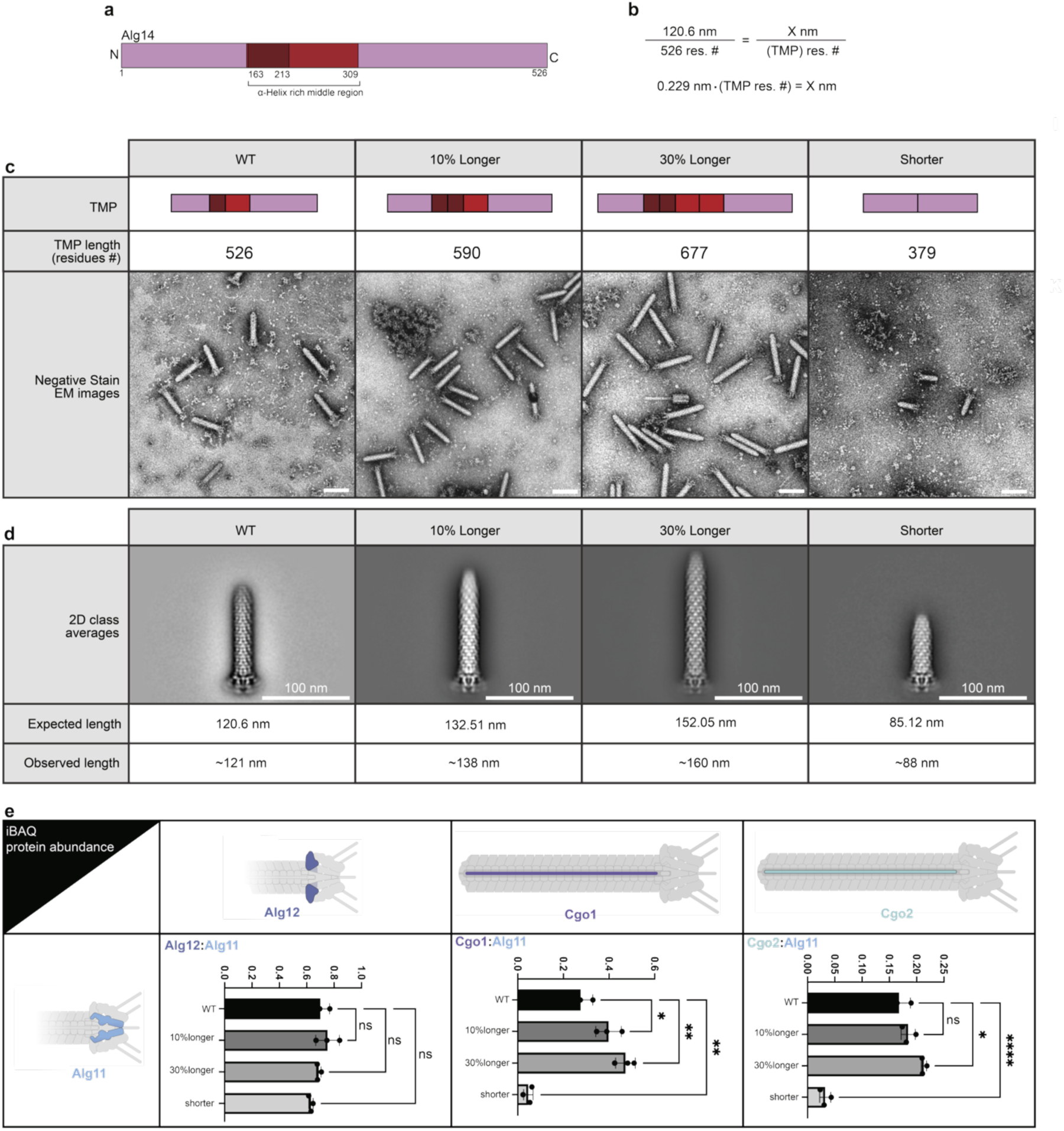
Tape measure protein controls AlgoCIS length and amount of loaded cargo. **a.** Diagram of the Alg14 tape measure protein. The internal α-helical region is shown in red, while the region containing the α-helices with the highest confidence scores from AlphaFold3 is indicated in dark red, the full AlphaFold3 structure of Alg14 is shown in Extended Data 5c. **b.** The ratio based on the length of WT AlgoCIS to the number of residues in Alg14 was used to identify a coefficient of 0.229, representing the nm of the final particle length per residue of TMP protein. **c.** Chart showing several Alg14 modification mutants, their residue length, and negative staining EM images of each corresponding mutant. Bars, 100 nm **d.** Representative 2D class averages of the negatively stained Alg14 mutant particles showed a descrete size and similar organization of the baseplate when compared to WT while the length of the sheath-tube module increased. The calculated expected length of the particles by multiplying the length of the TMP with the 0.229 nm residue coefficient. The observed length of the mutant Alg14 particles were determined from the major classes of the 2D projections, revealing that the observed lengths were relatively consistent **e.** The increased volume of the inner tube lumen allowed for larger amounts of cargo proteins to be loaded per AlgoCIS particle. MS iBAQ protein abundance ratios of purified AlgoCIS particles were used to investigate the ratios of AlgoCIS proteins to one another. The ratio of two standard baseplate proteins, Alg11 and Alg12 were used as a negative control due to the ratio between them staying consistent regardless of particle sheath-tube length. The ratio between both Cgo1 and Cgo2 with the baseplate protein Alg11 indicated that more cargo proteins are found per particle in both the Alg14-10%-longer as well as Alg14-30%-longer mutants. The Alg14-shorter mutant showed a significant decrease in the ratio of both Cgo1 and Cgo2 to Alg11.

To determine if the longer AlgoCIS particles resulted in more cargo protein being loaded into the now larger volume of the AlgoCIS tube lumen, we set up a quantitative MS experiment. We rationalized that the number of baseplate proteins would remain constant per eCIS while the number of sheath or cargo protein would be variable in the differing length mutants. Therefore, we began by validating the assay by analyzing the iBAQ protein abundance ratios of the baseplate protein Alg11 to the other major baseplate protein Alg12. Each tail should have six copies of each baseplate protein, regardless of overall tail length. The ratios of Alg11:Alg12 for the WT AlgoCIS, Alg14-10%longer, Alg14-30%longer, and Alg14-shorter mutants were consistent among the mutants (Fig. 4e). When the same strategy was applied to the ratio of Alg11:Cgo1 and Alg11:Cgo2, there was an increase in Cgo1 and Cgo2 in the AlgoCIS-Alg14-10%longer and AlgoCIS-Alg14-30%longer particles when compared to WT, and a decrease was seen in the AlgoCIS-Alg14-shorter sample (Fig. 4e).

In summary, we show that the TMP (Alg14) determines the length of the AlgoCIS sheath-tube module and that length modifications can be engineered by manipulating the length of Alg14. The resulting length differences impact the amount of cargo that is loaded into the inner tube.

### Specific protein delivery to mammalian and bacterial cells

Finally, our goal was to deliver cargo of choice to both mammalian and bacterial cells. In order to perform functional assays with HeLa cells, we combined 1) retargeting of AlgoCIS using the tail fiber towards CAR by Ad5 (AlgoCIS-Ad5) and 2) loading AlgoCIS with beta-lactamase BlaM (AlgoCIS-BlaM), resulting in AlgoCIS-Alg19ΔTip::Ad5 / ΔCgo1-2::Cgo2LS-BlaM (AlgoCIS-Ad5/BlaM) (Fig. 5a). Mutant cells were cultivated, followed by AlgoCIS purification and co-incubation of particles with adherent HeLa cells for 1 h at 37 °C with 5% CO2. After incubation, the HeLa cells were loaded with the FRET-based BlaM reporter CCF2-AM^38^. CCF2-AM readily enters cells and undergoes a FRET emission shift from ∼535 nm (green) to ∼450 nm (blue) in the presence of active BlaMs^39,40^ (Fig. 5a). While we observed blue signals in the AlgoCIS-BlaM treated cells, a higher level of cleaved signal was observed in the AlgoCIS-Ad5/BlaM treated cells (Fig. 5b/c). Interestingly, there was a correlation between the number of cells observed to exhibit a blue color and the amount of AlgoCIS-Ad5/BlaM co-incubated with the HeLa cells (Fig. 5c). This indicated that engineered AlgoCIS injected functional BlaM protein into HeLa cells in a dose responsive manner (Extended Data Fig. 6).

**Fig. 5:**
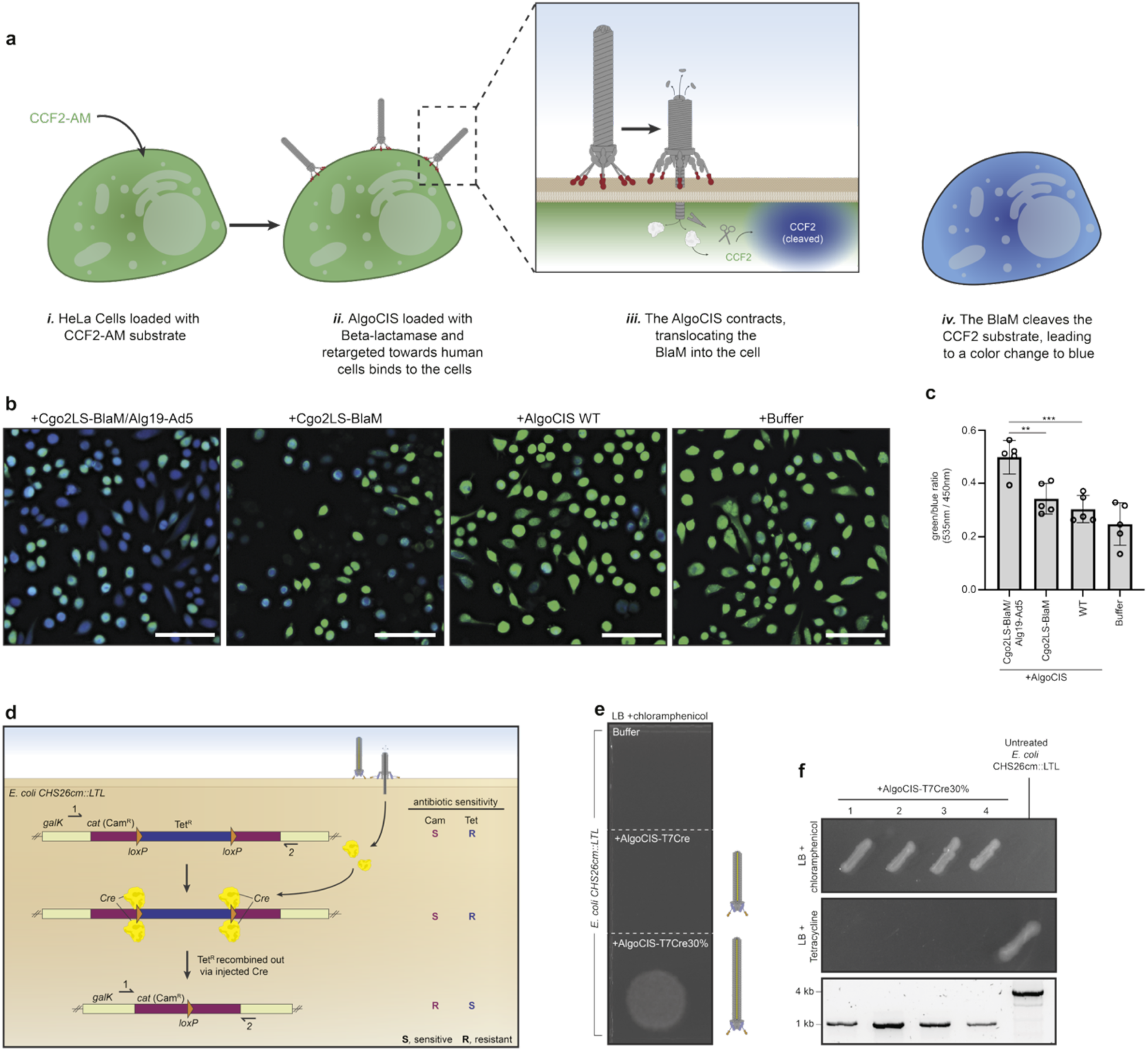
Combining retargeting, custom cargo loading, and length modification. **a.** Beta-lactamase (BlaM) translocation assay workflow. HeLa cells are loaded with the BlaM responsive substrate CCF2-AM. If BlaM is translocated through the cell membrane via the AlgoCIS-Ad5/BlaM mutant, the CCF2 beta-lactam ring will be cleaved, resulting in a FRET shift downwards from the uncleaved green (∼535 nm) to cleaved blue (∼450 nm). **b.** Representative qualitative LM images of CCF2-loaded HeLa cells post co-incubation with purified AlgoCIS. We observed an increase in cleaved signal (blue signal) in the AlgoCIS-Ad5/BlaM treated sample compared to the AlgoCIS-BlaM, WT, and buffer treated samples. Bars, 100 µm. **c.** A bar graph showing the quantification of CCF2 signal in each cell, calculated by the ratio of green (535 nm) to blue (450 nm) signal in each cell. Images were quantified per cell, using ImageJ, with each point representing a technical replicate consisting of >300 cells. **d.** Schematic of the Craft assay utilizing the chromosomal *loxP* reporter *E. coli* strain CHS26cm::LTL. The non-essential *galK* gene is interrupted by a chloramphenicol resistance cassette, which is in turn interrupted by a tetracycline resistance cassette that is flanked by *loxP* sites. When Cre is present, the two *loxP* sites that are in parallel will recombine out the Tet^R^ cassette, functionally flipping the antibiotic resistance of the *E. coli* strain from Tet^R^ to Cam^R^. Primers 1 and 2 are used to confirm the Cre mediated recombination. The antibiotic resistance of each strain is shown on the right. **e.** Post AlgoCIS treatment, the *E. coli* strains were grown overnight in LB, then serially diluted and dropped onto an LB plate containing chloramphenicol to evaluate recombination. Only the longer and retargeted version showed any colonies, indicating that the Cre was able to recombine the *loxP* sites flanking the Tet^R^ gene, resulting in the *cat* gene being expressed and *E. coli* becoming resistant to chloramphenicol. **f.** Colonies of treated *E. coli* colonies were confirmed to have recombined out the Tet^R^ gene with a colony PCR. Primers 1 and 2 shown in panel d were used to confirm the recombination of the loxP, resulting in a smaller PCR product on positive colonies.

For injection into bacterial cells, we generated mutants that were retargeted towards *E. coli* via the T7 tail fiber tip replacement and loaded with the reporter protein Cre. Purified AlgoCISs were incubated with a Cre reporter strain, *E. coli* CHS26cm::LTL for 3 h at 37 °C shaking at 200 RPM. The reporter strain contains a floxed tetracycline resistance cassette (Tet^R^) interrupting a chloramphenicol resistance cassette (Cam^R^) (Fig. 5d/e). Post incubation, the *E. coli* culture was plated onto LB media containing chloramphenicol as a selective marker for Cre-mediated recombination (Fig. 5f); however, no *E. coli* colonies formed from samples treated with AlgoCIS-T7/Cre particles. We reasoned therefore that either insufficient cargo amount or insufficient particle length may prevent successful injection into the cytoplasm. We thus further modified the particles by making them 30% longer as described above (AlgoCIS-T7/Cre/30%), resulting in the appearance of colonies. To confirm that successful injection took place, we patch plated the colonies in replicates onto chloramphenicol plates as well as on tetracycline containing plates. These colonies were then further tested via PCR, utilizing primers flanking the *loxP* recombination region, with the PCR products showing sizes indicative of Cre mediated *loxP* recombination (Fig. 5g). Sequencing of the PCR products verified successful recombination resulting from the AlgoCIS-T7/Cre/30% treatment.

## Discussion

Contractile injection systems are a powerful emerging biotechnological tool, featuring key advantages such as the coupling of cell type-specific targeting paired with custom protein payload delivery. While the potential of this family of systems is becoming apparent, insights into their molecular mechanisms remain scarce. Here we establish PROCIS as i) a mechanistic framework for eCIS research and ii) a biotechnological tool for protein delivery (summarized in Fig. 6). The addressed mechanistic aspects include target binding, cargo loading and length determination, which are discussed below.

**Fig. 6:**
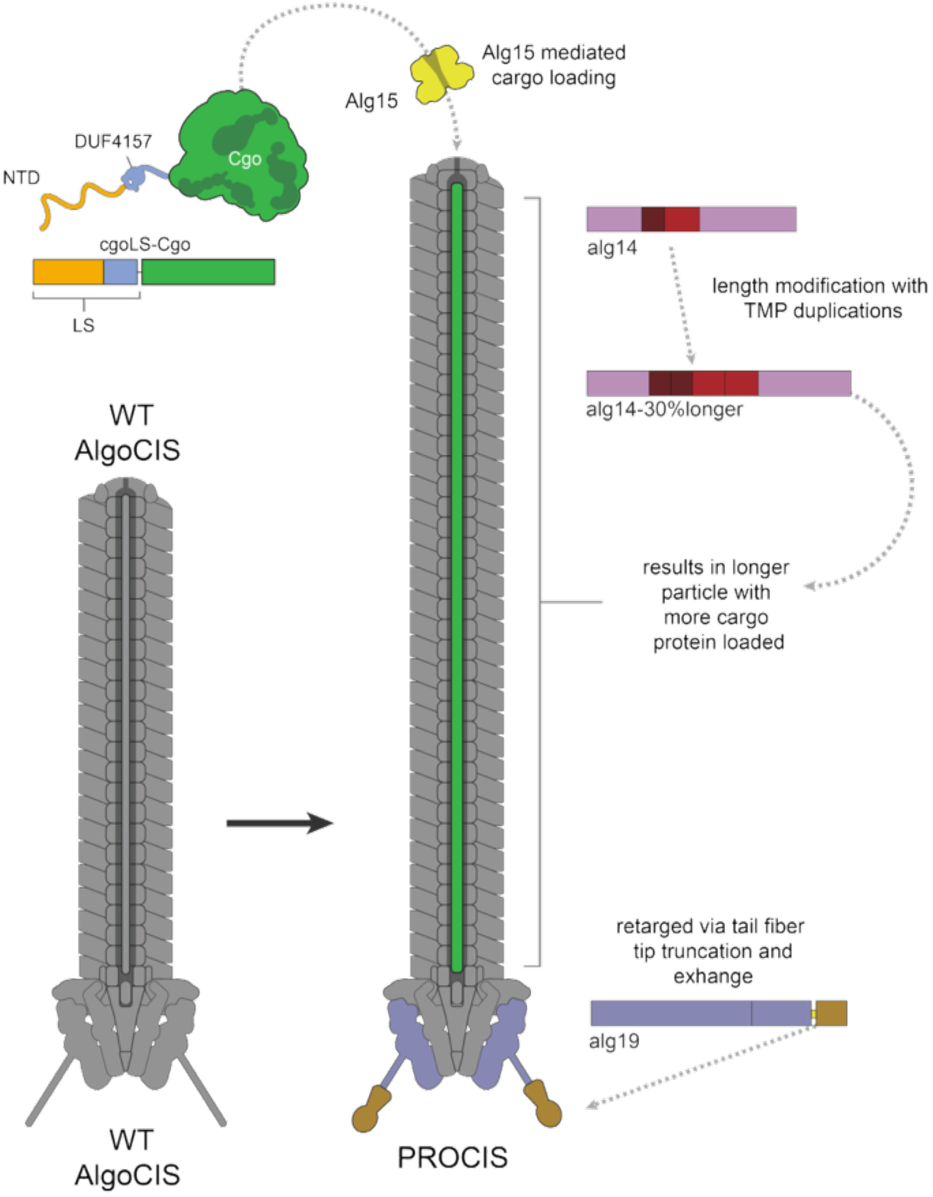
Overview of PROCIS as a modular platform for protein delivery. PROCIS can be re-engineered in three ways, 1) loading custom cargo into the inner tube lumen which is mediated via the Alg15 AAA+ protein, 2) retargeting the CIS towards novel targets via replacing the tip of the tail fibers, and 3) length change of the sheath-tube module by lengthening the TMP, resulting in more cargo being loaded.

### Target binding

Tail fibers were shown previously to determine the specificity of phages, pyocins and eCIS to bind to specific target cells. The replacement of (partial) tail fibers was reported as a strategy to alter target specificity^15,41,42^. The sequences, and molecular structures of tail fibers, however, are highly variable and not conserved even across closely related systems. We therefore employed structure prediction to identify the precise C-terminal domain that could be exchanged with domains that allowed for binding target cells of choice. Interestingly, the same truncation of the tail fiber combined with three different binding modules turned out to result in successful binding to target cells and subsequent firing into bacterial and mammalian cells. Together with the diverse range of targeted cell types (wildtype bacteria, engineered surface structures, and mammalian cells), this highlights the robustness and adaptability of the system. This observation is also consistent with our finding that Alg19-like proteins reveal a conserved domain structure and show highest divergence in the flexible C-terminal fiber-like domain (Extended Data Fig. 7).

### Cargo loading

An N-terminal ‘packing domain’ has been identified in cargo proteins that are loaded into PVCs; however, the mechanism of loading remains unclear for any eCIS. Based on our data, we propose a model for the loading of proteinaceous cargo into the lumen of the inner tube (Fig. 6). We show that an N-terminal cargo loading sequence acts as a signal sequence that directs native and non-native cargo into the tube lumen. This process is mediated by the AAA+ protein Alg15, which likely unfolds the cargo for loading. Downstream of the N-terminal LS, some but not all eCIS cargo proteins encode a DUF4157 domain, a domain that has been correlated with potential eCIS effector proteins^28,43^. Our results indicate that DUF4157 is not critical for loading (cargo without DUF4157 is still loaded; Fig. 2d) and it may therefore play a dispensable/accessory role for loading or a role that is associated with the respective cargo protein (e.g. refolding in the target). By analyzing the N-terminal LS of cargo proteins of different eCIS, we find that it is not conserved. Sequence analyses however show a clear trend that indicates a high degree of intrinsic disorder in this LS (Extended Data Fig 7). We hypothesize that the LS may be specific for and recognized by the respective AAA+ protein.

AAA+ proteins play a wide range of roles across domains of life by their ability to unfold proteins^44,45^. As typically ring-shaped, hexameric protomers, they generally unfold substrates by threading them through a central channel. We determine Alg15 as a critical AAA+ factor for loading cargo into the tube lumen and hypothesize that this process may either take place during the assembly of the AlgoCIS particle or after structural assembly has been completed. It is likely that the AAA+ protein dissociates from the AlgoCIS particle after loading, since Alg15 has not been detected by either proteomics or cryo-EM of fully assembled and loaded particles. The unfolded nature of cargo inside the tube lumen is supported by the large size of cargo proteins that can be loaded and the absence of structured domains in the lumen in cryo-EM maps of loaded particles (Fig. 2e/f/g). Furthermore, this hypothesis is supported by the inability to load stably folded proteins such as sfGFP (Fig. 2f), which Alg15 may not be able to unfold. This finding also informs future engineering projects, identifies protein stability as a critical cargo characteristic, and it also poses the question whether stronger AAA+ proteins may be engineered that could overcome this limitation.

### Length determination

Finally, we identified a strategy to control the length of engineered eCIS particles and therefore also the amount of loaded cargo. This was accomplished by engineering the TMP, which controls the length of bacteriophage tails, eCIS and some T6SSs^29,34,46^. The TMP is thought to run through the lumen of the inner tube, with the length of their protein sequence correlating to the length of the contractile particle. In many cases, the TMP remains inside the inner tube lumen of the assembled particle^47,48^. Since we could not detect the TMP by MS of purified AlgoCIS, we suggest that the TMP is dissociated prior to the completion of assembly. This may be due to an evolutionary maximization of cargo load per particle. In fact, our data reveal that the length of the eCIS particle is a critical factor that determines the amount of cargo loaded per eCIS particle, with implications on future engineering of eCIS. Since TMPs are generally conserved across different systems, the approach shown here will be applicable to other eCIS in the future.

### Integration of results for protein delivery to bacterial and mammalian cells

The integration of the above insights with functional assays allowed us to test the delivery of non-native cargo to bacterial and mammalian cells. The data indicate that engineered AlgoCIS remained contractile, assumed the ability to bind to new targets and accommodated non-native cargo that was functional and active in the target organism. The failure of engineered AlgoCIS with WT length to induce a cellular response in bacteria presents an exciting avenue for future research, which will examine whether longer AlgoCIS versions were successful because of their higher load of cargo or because of their longer reach that may be needed to penetrate through the cell envelope and periplasm of the bacteria.

In conclusion, we demonstrate that eCIS are highly modular and amenable to modifications that confer new functionalities. By integrating retargeting, custom cargo loading, and length adjustments, we show that PROCIS can deliver active proteins into both bacterial and mammalian cells, highlighting the platform’s capacity to accommodate multiple engineered features simultaneously. Furthermore, our study will serve as a framework for research into the molecular mechanisms of eCIS.

### Statement of contributions

CFE and MP conceived the project; CFE/ET/DS designed and generated all mutant strains; CFE/ET/DS purified samples; CFE performed killing assays on both eukaryotic and bacterial cultures and analyzed the data; CFE/DS prepared cryo-EM/ET samples, collected cryo-ET data, and analyzed the reconstructed tomograms; CFE/DS prepared, collected, and analyzed the light microscopy data; PA/CFE/DS collected and processed cryo-EM data, built and refined structural models, and performed structural analysis; PA performed SPA of negatively stained samples. CFE and MP wrote the manuscript with input from all authors.

## Supporting information

SuppTable1_StrainPlasmid_Table

SuppTable2_MS_Summary

## Acknowledgements

We thank Jingwei Xu for providing scientific feedback on the experimental design and for support with cryo-EM data collection and analysis. We are also grateful to Ellen Zechner for generously providing the *E. coli* CHS26cm::LTL strain. We thank ScopeM for instrument access at ETH Zürich. We thank the Functional and Genomic Center Zürich (FGCZ) for mass spectrometry support. Pilhofer Lab members are acknowledged for discussions. M.P. was supported by the Swiss National Science Foundation (31003A_179255 and 310030_212592), the European Research Council (679209 and 101000232), and the NOMIS foundation.

## Materials and Methods

### Bacterial growth conditions and generation of mutants

All *A. machipongonesis* PR1 cultures were grown in <u>m</u>arine <u>b</u>roth (MB) (Condalab) at 30 °C and 200 RPM shaking and with 1.5 % agar when grown on plates unless otherwise stated. For the generation of mutants in *A. machipongonesis* PR1, the previously established protocol was followed^12^. Briefly, the pCHIP3 plasmid was used to integrate into the *A. machipongonesis* genome via homologous recombination with successful integration selected for by selection on MB plates containing 50 µg ml^—1^ of erythromycin. Colonies with successful plasmid integration were grown in 25 mL MB broth then plated on MB containing 10 mM of 4-chloro-DL-phenylalanie and grown at 30 °C for at least 72 hrs for counter-selection. Resulting individual colonies were screened for correct homologous recombination via colony PCR on both the counter-selection and selection plates. All strains and plasmids used in this study are summarized in Supplementary Table 1.

### AlgoCIS sheath preparation

The AlgoCIS purification was followed as previously reported. Briefly, a small volume of 25 mL of *A. machipongonesis* PR1 was inoculated into 1 L of fresh MB medium and grown at 30 °C and 200 RPM for 48 hrs. The bacterial pellet was harvested by centrifugation at 7000 rcf for 20 mins then resuspended with ∼20 mL MB medium. The lysis reagents (1% Triton-X100, 0.5× CellLytic B (Sigma-Aldrich), 200 μg/mL lysozyme, and 50 μg/mL DNAse I were added into the bacterial suspension and incubated at 37 °C for 30 mins to lyse the bacteria. The cell debris were removed by centrifugation at 21,000 rcf and 4 °C for 20 mins. The supernatant was subjected to ultra-centrifugation with sucrose cushion (1 mL at bottom) (20 mM Tris pH 8.0, 150 mM NaCl, 50 mM EDTA, 1% Triton-X100, 50% {w/v}sucrose) at 150,000 rcf and 4 °C for 1 hr. The sucrose cushion was taken, together with some remaining overlying liquid (∼ 0.5 mL). The residual bacterial contamination in solution was further removed by centrifugation at 21,000 rcf for 20 mins. The sample was subjected to a second round of ultra-centrifugation without a sucrose cushion. The resulting pellets were washed, soaked with buffer A (20 mM Tris pH 8.0, 150 mM NaCl, 50 mM EDTA) overnight, and then resuspended. The sample was further purified through a 10–50% (w/v) sucrose gradient at 100,000 rcf and 4 °C for 1 hr using the SW 55 Ti rotor. The gradient was fractionated into 11 fractions and fractions 4-9 were collected. The solution was then diluted with HEPES buffer (20 mM Tris pH 7.5 and 150 mM NaCl), and passed through a 0.2-μm-pore filter twice, then concentrated by a third round of ultra-centrifugation at 150,000 rcf for 1 hr. The pellets were resuspended in buffer B and stored at 4 °C. The purified samples were analyzed by negative staining and EM and the concentration was quantified by A280 measurement on a Nanodrop instrument (ThermoFisher).

### Co-incubation of AlgoCIS with target cells

For *E. coli* strains, the cultures were grown in at 37 °C, 200 RPM LB with the appropriate antibiotics added when necessary (100 µg/mL ampicillin, 50 µg/mL kanamycin, 10 µg/mL tetracycline, 25 µg/mL chloramphenicol). The *E. coli* strains were grown overnight before being diluted to an OD600 of 0.1 in LB. Purified AlgoCIS were normalized to 1.0 mg/mL before being co-incubated with the cells overnight at 37 °C, 200 RPM. The cells were plated on LB agar square plates in 1:10 serial dilutions before being incubated at 37 °C overnight.

The following day the plates were taken out and were imaged via the colorimetric setting on a gel imager.

For HeLa cell i*n vitro* co-incubation assays, HeLa cells were grown in multi-well dishes to a confluency of around 70–80% at 37 °C and 5% CO2, with *n* = 4 for each assay. Purified AlgoCIS were normalized to 1.0 mg/mL with the eukaryotic cell growth media. The medium in the multi-well dish was removed and replaced with the AlgoCIS-containing media and the plate was incubated back at growth temperatures. The plates were observed for change at 24 hr, 48 hr and 72 hr post treatments. Cells were live/dead stained with FDA/PI and images were taken of each well via fluorescent light microscopy (see below for parameters).

### Immunofluorescence and Light microscopy

For immunofluorescence assays, the cultures were fixed with 4% of prewarmed paraformaldehyde at 37 °C for 15 mins. The samples were then washed with 1.5 volumes of PBS three times. The samples were then blocked with a 3% bovine serum albumin (BSA) in PBS solution while rocking at room temperature for 1 hr. The samples were then again washed again with 1.5 volumes of PBS three times. The primary antibody solution of 1:100 dilution of rabbit α-Alg2 (anti-sheath) in 0.1 % bovine serum albumin solution was added and incubated rocking at room temperature for 1 hr. The samples were washed again before a secondary solution of 1:500 dilution of FluoTag-X4 anti-rabbit IgG (NanoTag Biotechnologies) and 10 µg/mL of Hoechst 33342 (Thermo Fisher Scientific) in 0.1 % bovine serum albumin solution protected from light and incubated at room temperature for 1 hr. The samples were washed in 1.5 volumes of PBS three times before being mounted with Vectashield (Vector Laboratories, for eukaryotic samples) or spotted on an 1.5 % low melt agarose pad (for bacterial samples) and imaged. Light microscopy data was collected on a Leica THUNDER wide field microscope and a Zeiss LSM900 Airyscan2 confocal microscope. CCF2-loaded cells were quantified using ImageJ software (National Institutes of Health, USA).

### Minicell Preparation

*E. coli* Δ*minCDE*, *mreB*-A125V was grown in LB at 37 °C 200 RPM overnight. The culture was then centrifuged at 5000 rcf for 20 mins at 4 °C. The supernatant was then taken and moved to 2 mL tubes being careful to not disturb the pellet. The 2 mL tubes were then centrifuged at 21,000 rcf for 40 mins at 4 °C. The resulting small pellets were carefully resuspended in LB media before being pooled together and centrifuged again at 21,000 rcf for 40 mins. The resulting pellet was again resuspended in LB with the supernatants being discarded. The purified minicells were then used for binding assays and EM analysis.

### Mass spectrum analysis

The purified AlgoCIS samples were sent in solutions to the Functional Genomics Center Zürich (FGCZ), which performed the mass spectrum and the subsequent data analysis. The samples were first digested by trypsin. These digested samples were dried and dissolved in 20 μL ddH2O with 0.1% formic acid. The samples were transferred to autosampler vials for liquid chromatography–mass spectrometry analysis (LC–MS/MS). The samples were diluted at the ratio 1:40, with 1 μL of each sample being injected on a nanoAcquity UPLC coupled to a Q-Exactive mass spectrometer (ThermoFisher).

The acquired MS data were converted to a Mascot Generic File format and were processed for identification using the Mascot search engine (Matrixscience). In addition, the acquired MS data were imported into PEAKS Studio (Bioinformatic Solutions) and were searched against the *Algoriphagus machipongonensis* PR1 database. The results were visualized by Scaffold software. For the iBAQ values, the acquired MS data were processed using DIANN^49^. Spectra were searched using the Uniprot *Algoriphagus machipongonensis* PR1 database (Taxon ID 388413) in FASTA file, and common protein contaminants. Carbamidomethylation of cysteine was fixed modifications, while methionine oxidation was variable. Enzyme specificity was set to trypsin/P, allowing a minimal peptide length of 7 amino acids and a maximum of two missed cleavages. The R package prolfqua^50^ as used to analyze the differential expression and to determine group differences, confidence intervals, and false discovery rates for all quantifiable proteins^50^. Starting with the report.tsv file generated by DIANN, which does report the precursor ion abundances for each raw file, we determined protein abundances by first aggregating the precursor abundances to peptidoform abundances. Then, we employed the Tukeys-Median Polish to estimate protein abundances. Furthermore, before fitting the linear models, we transformed the protein abundances using the variance stabilizing normalization^51^. Calculating the iBAQ signal per protein involves summing the Precursor. Quantity values across all precursors associated with that protein and dividing by the number of theoretically observable peptides of the protein. Summarized MS data on AlgoCIS can be found in Supplementary Table 2.

### Vitrified sample preparations

The purified AlgoCIS particles were vitrified on 200 mesh Quantifoil Gold grids (R 2/1) using a Vitrobot Mark IV (Thermo Fisher Scientific). For cryo-ET, different samples were seeded with 10 nm BSA-coated colloidal gold particles at the ratio 1:5 before application to EM grids. For tomography, grids were glow discharged at 25 mA for 45 sec before 3 µL of sample application. For single-particle analysis, an additional 1 nm of carbon was added to the surface of the grids.

### Single-particle cryoEM analysis

Cryo-EM datasets of purified AlgoCIS were collected on Titan Krios microscope (Thermo Fisher Scientific), operated at 300 keV equipped with K3 direct electron detector, operating in counting mode, and using a slit width of 20 eV on a GIF-Quantum energy filter (Gatan). The automated SPA collection was conducted with EPU software (Thermo Fisher Scientific) at 81000x magnification with a final pixel size of 1.06 Å/pix over 50 frames with a total dose of 70 e^-^/Å^2^. The targeted defocus was set between -1.8 and -3 µm with 0.2 µm increment. Data collection was monitored by in-house *data_collection_alarm.py* program^52^.

Single-particle analysis was performed in CryoSPARC v4.5.3^53^ using approach described in Casu et al, 2023^54^.The gain reference was estimated *a posteriori*^55^ and used in movie-alignments, followed by CTF-determination and automatic particle picking using Filament tracer. For the ΔAlg15 and WT filled vs empty dataset, only end-particles of the filaments very selected from the corresponding 2D-classes, and 1,366 baseplate-containing particles were processed separately from 910 cap-containing particles. *Ab initio* created models were built and used in 3D-refinements (Homo Refine) with applied C6-symmetry.

Cryo-ET datasets of samples were collected at a nominal magnification of 19’500x with a pixel size of 4.5106 Å/pix and a dose-symmetric scheme from -61° to 61° at 2° incremental steps using SerialEM. The total dose rate of each collection was ∼123e^−^/Å^2^. The tomograms were aligned and reconstructed with the IMOD^56^ program suite.

### Negative staining EM dataset collection

Negative staining EM datasets for the length determination were collected on the Talos L120C microscope (Thermo Fisher Scientific), equipped with Ceta S 16M CMOS detector, operating in automatic mode with the EPU v3.4 software. The data was collected at a magnification of 28,000x (pixel size 5.01 Å/pix) with a total dose of about 15 e^-^/Å^2^. The target defocus was set to -2 μm. 747 micrographs were collected for the Alg14-shorter dataset and 386 for the Alg14-30%longer. Both datasets were processed in CryoSPARC v.4.5.3 software without CTF correction. Manually selected particles (24 for the Alg14-shorter and 72 for the Alg14-30%longer dataset) were subjected to 2D-classification with five classes to create templates for correlation particle picking. Picked particles were subjected to several rounds of 2D-classification to reveal discrete 2D class-averages corresponding to particles with well-defined lengths.

### *In silico* characterization of cargo proteins

Proteins containing the DUF4157 domain were identified via InterPro^57^ and subsequently processed using pfamScan^58,59^ to define domain architectures and precise residue coordinates. Each sequence was partitioned into three distinct segments based on these coordinates: the N-terminal domain (NTD), the DUF4157 core, and the C-terminal domain (CTD). To assess regional protein flexibility, the intrinsic disorder of each segment was predicted using IUPred2A^60^ (10.4.1).

Root-mean-square fluctuation (RMSF) values for the cargo proteins were calculated using the CABS-flex 2.0 web server^61^. Phylogenetic trees of the cargo NTDs were calculated and visualized using the MegaX software^62^.

---

**Extended Data Fig. 1:**
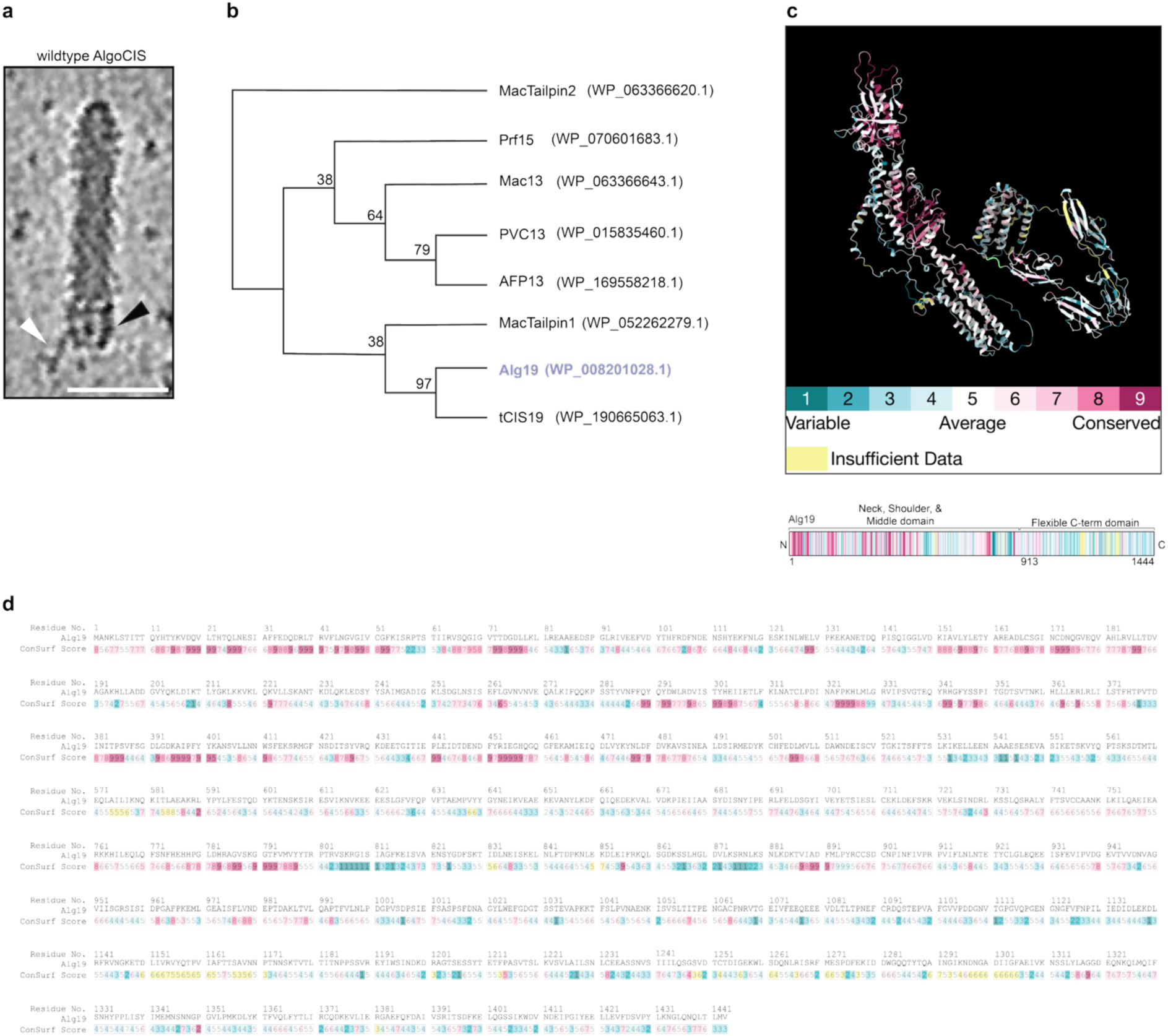
Tomographic visualization, phylogeny, and conservation mapping of the Alg19 tail fiber. **a.** Tomographic slice of an AlgoCIS, black arrowhead indicates the shoulder, neck, and middle domains of the Alg19 tail fiber. The white arrowhead indicates one of the flexible C-terminal domains. Bar, 50 nm. **b.** A phylogentic tree of known eCIS-like tail fiber proteins. Alg19 and tCIS19 group separately from previously studied CIS tail fiber proteins. Numbers represent bootstrap values when run over 1’000 iterations in the MegaX software. **c.** ConSurf analysis mapped amino acid conservation onto the Alg19 monomer structure. The C-terminal flexible domain exhibits low conservation relative to the structured shoulder, neck, and middle domains. **d.** ConSurf score values shown for each residue of the Alg19 protein. Residues are colored with the same key as panel c.

**Extended Data Fig. 2:**
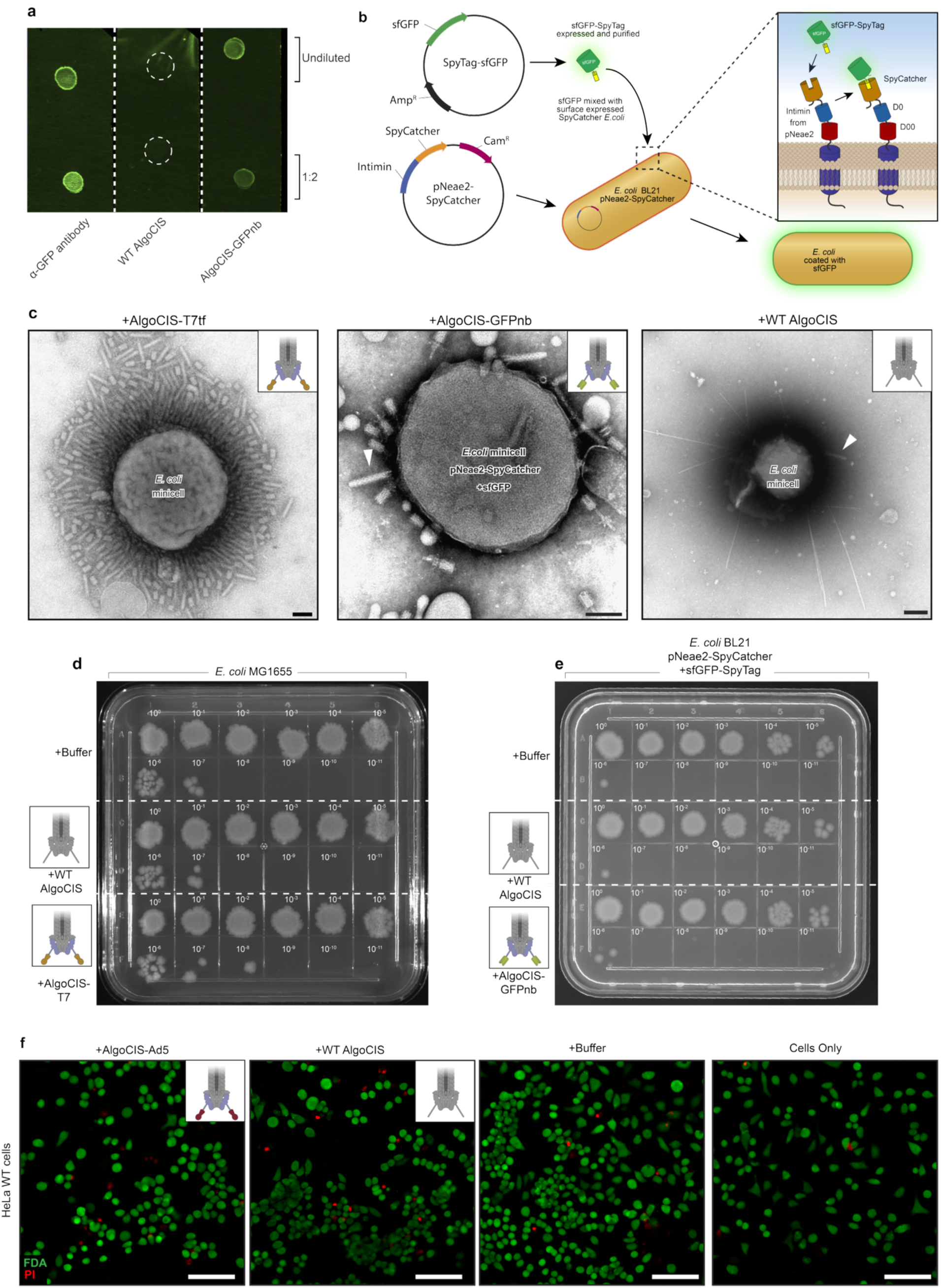
Functional validation and targeting assays of engineered AlgoCIS variants. **a.** A dot blot assay to validate the AlgoCIS-GFPnb mutant for its ability to bind free floating GFP. WT AlgoCIS, AlgoCIS-GFPnb, and an α-GFP antibody (positive control) were spotted on a nitrocellulose membrane and allowed to dry. The rest of the membrane was then blocked with 3% BSA before being washed with purified sfGFP before being washed and imaged on a fluorescent gel imager. The AlgoCIS-GFPnb and α-GFP antibody spots showed GFP signal where the samples had been applied, while the WT AlgoCIS exhibited no signal. **b.** Workflow schematic for coating *E. coli* with sfGFP. The intimin based surface expression plasmid pNeae2 was fused with SpyCatcher in order to express it on the bacterial surface. Exogenously purified sfGFP tagged with SpyTag was incubated with an *E. coli* culture which was induced to express the intimin-SpyCatcher fusion. The cells were then washed before being co-incubated with AlgoCIS samples and analyzed. **c.** Negative Stain EM images of *E. coli* minicells after co-incubation with AlgoCIS-T7 (left), minicells coated with sfGFP then co-incubated with AlgoCIS-GFPnb (middle), or *E. coli* minicells WT AlgoCIS control samples. Retargeted AlgoCIS were seen frequently bound to the minicells while the WT AlgoCIS control sample rarely had AlgoCIS associated with a minicell. Example AlgoCIS are indicated with white arrowheads in the AlgoCIS-GFPnb and WT control images. Bars, 100 nm. **d-e**. Killing assay of the AlgoCIS-T7 after co-incubation with *E. coli* MG1655 and *E. coli* BL21 coated with sfGFP co-incubated with AlgoCIS-GFPnb respectively. The bacteria were co-incubated with purified AlgoCIS for 4 h rocking at room temperature before being 10-fold serially diluted and spotted onto an LB plate, and grown at 37 °C overnight before being imaged. **f.** Representative images of HeLa cell killing assays. HeLa cells co-incubated with AlgoCIS for 24 h before being washed and stained with fluorescein diacetate (FDA, green) as a live stain and propidium iodide (PI, red) as a dead stain. There was no discernible difference between the retargeted AlgoCIS-Ad5, WT AlgoCIS, or buffer control samples. Bars, 100 µm.

**Extended Data Fig. 3:**
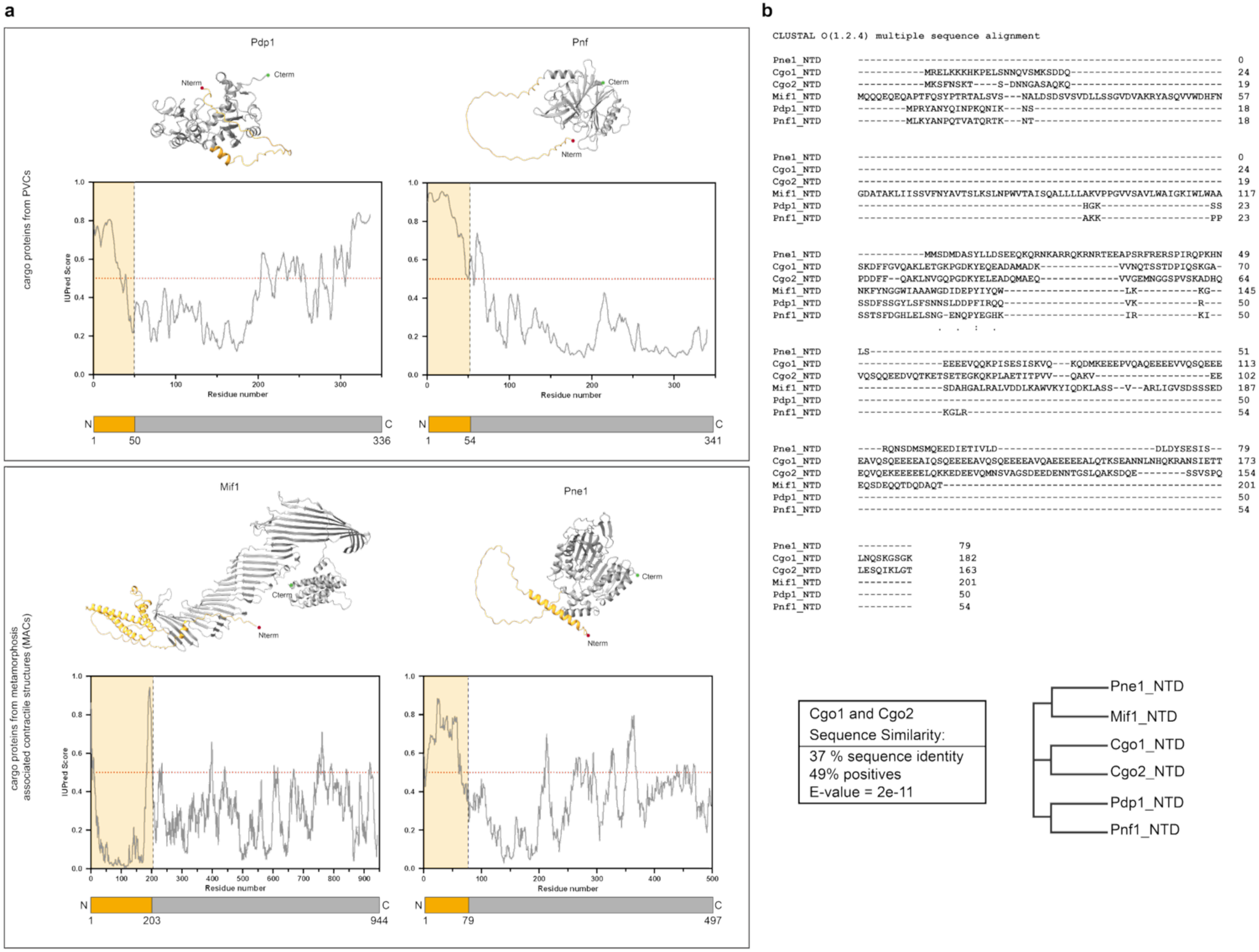
Other eCIS cargo proteins contain a disordered NTD. **a.** AlphaFold3 models for the cargo proteins from related eCIS systems with the IUPred2a graphs directly below the structures. The two Photorhabdus virulence cassette (PVCs) cargo proteins Pdp1 and Pnf both contain NTD “packaging domains,” but notably do not consist of a predicted DUF4157. The two Metamorphosis associated contractile structures (MACs) cargo effectors Mif1 and Pne1 vary in their N-terminal region, with Mif1 only While Pne1 has a more clearly disordered NTD, the Mif1 cargo protein has a less well defined disordered NTD. **b.** Clustal Omega alignment of the NTDs from the Cgo proteins from AlgoCIS (Cgo1 and Cgo2), PVC (Pdp1 and Pnf1), and MACs (Mif1 and Pne1), along with the alignment statistics from the Cgo1 and Cgo2 alignments. The phylogenetic tree of the NTDs show that they cluster based on the host bacteria.

**Extended Data Fig. 4:**
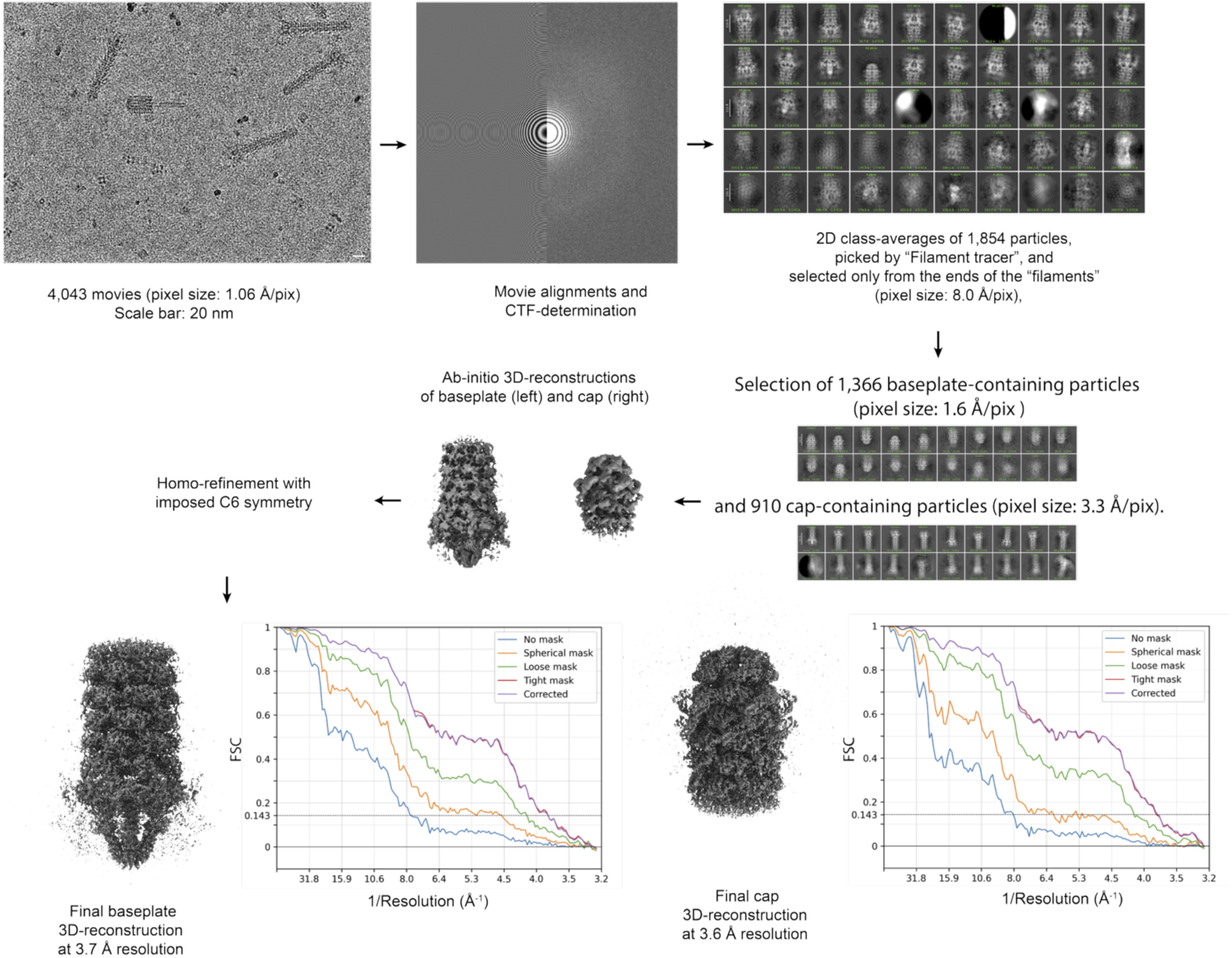
Cryo-EM SPA workflow. Flowchart for the cryo-EM single particle reconstruction of the AlgoCISΔAlg15 baseplate and cap portion to obtain the structures shown in Figure 2D.

**Extended Data Fig. 5:**
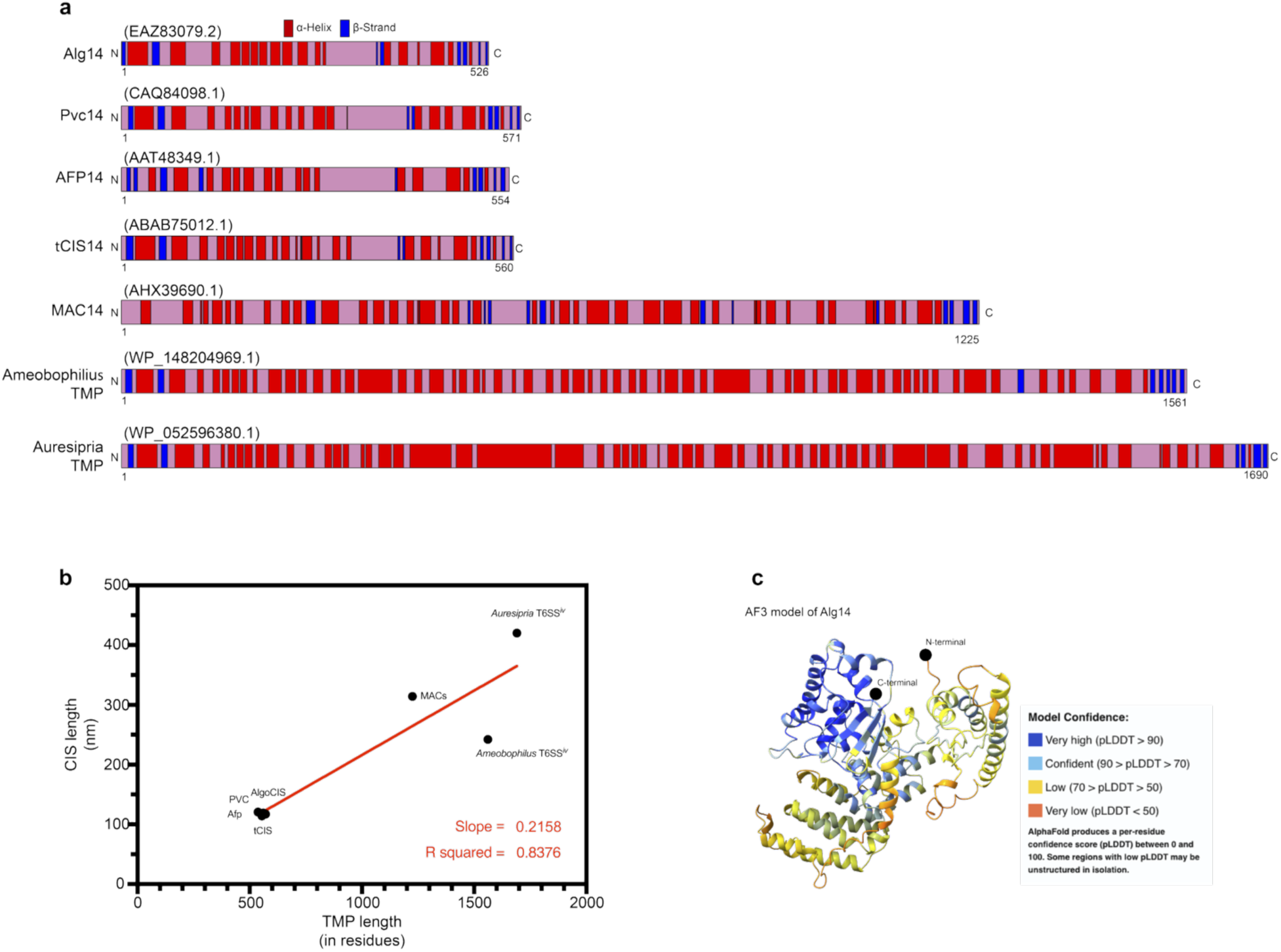
eCIS tape measure proteins contain a central region which is predicted to be comprised of mainly α-helices. **a.** The tape measure proteins of the known eCIS-related particles with their predicted secondary structures shown. Secondary structures were predicted with the Phyre2 secondary structure prediction tool, red corresponds to predicted α-helices while blue indicates predicted β-strands. **b.** The tail lengths of selected CISs plotted against the length (amino acid residues) of their corresponding tape measure proteins. Red line represents best fit line and has a slope of 0.2156 and R-squared value of 0.8376. **c.** The AlphaFold3 prediction of Alg14, colored with the pLDDT confidence scores. The highest confidence α-helices (seen in top left bundle) were chosen for modification. N- and C-termini are shown with black circles.

**Extended Data Fig. 6:**
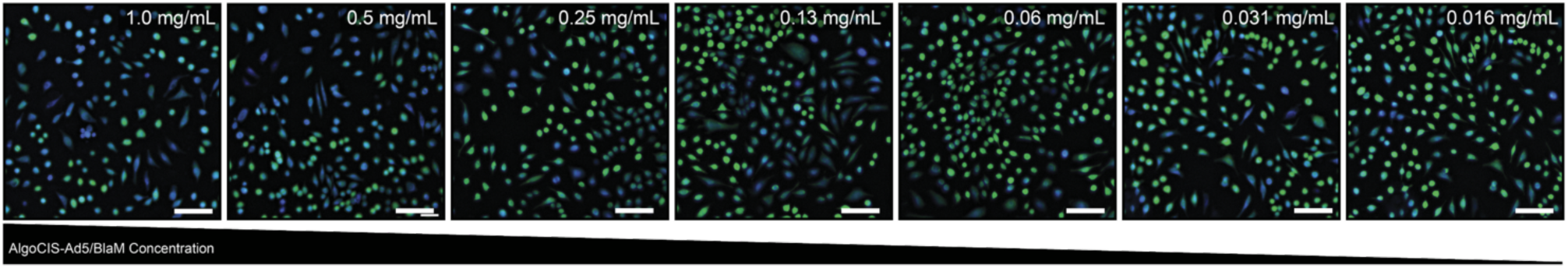
Dose response of AlgoCIS-mediated BlaM injections into HeLa cells. Qualitative images of functional assays to test AlgoCIS-Ad5/BlaM-mediated injection into HeLa cells (loaded with CCF2-AM) with decreasing concentrations of AlgoCIS added. Higher concentrations of AlgoCIS correlate with higher levels of blue signal. Bars, 100 µm.

**Extended Data Fig. 7:**
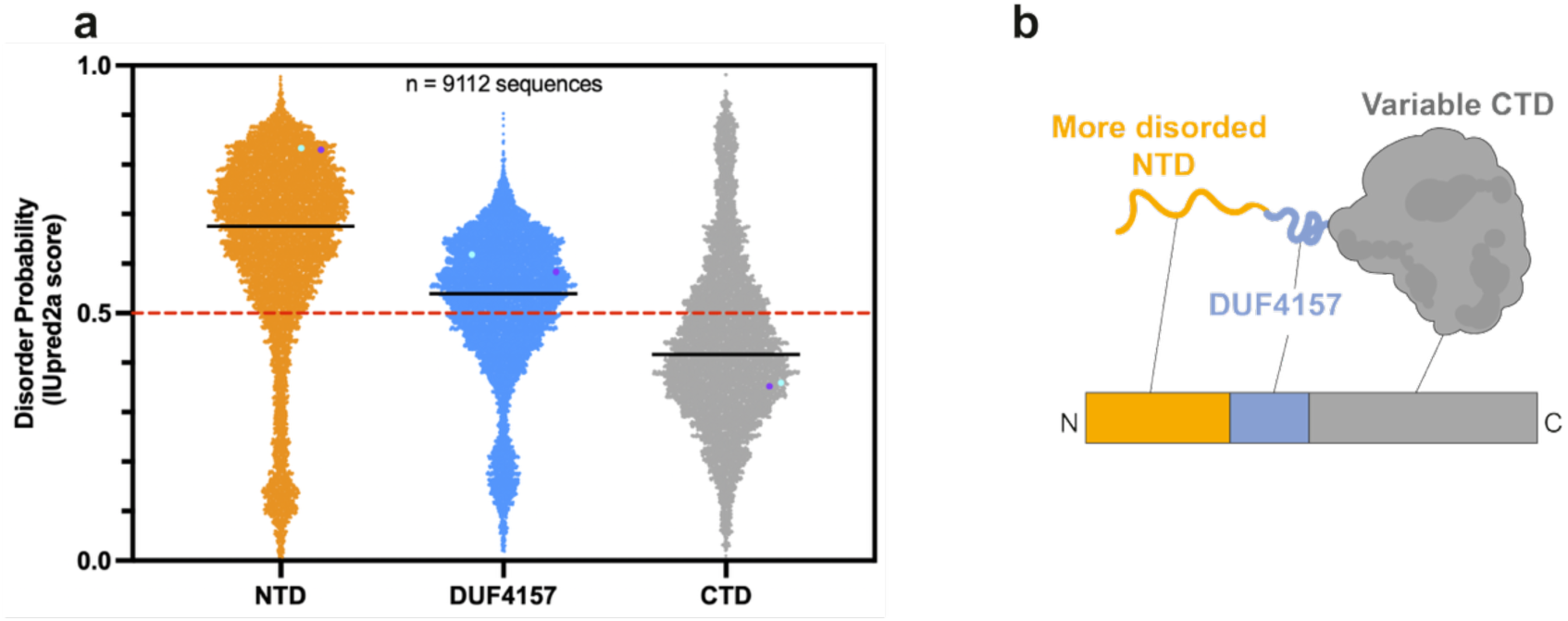
*In silico* disorder prediction of DUF4157 containing proteins. **a.** Protein sequences containing the predicted DUF4157 domain were retrieved from InterPro. Sequences were partitioned into three distinct regions based on domain architecture: the N-terminal domain (NTD), the DUF4157 domain, and the C-terminal domain (CTD). The intrinsic disorder of each region was independently quantified using IUPred2a. Individual data points represent domain-specific disorder scores, with Cgo1 (purple) and Cgo2 (teal) highlighted for reference. These results demonstrate that the NTD possesses a higher predicted disorder propensity compared to the relatively ordered DUF4157 and CTD regions. **b.** Proposed cartoon model of a DUF4157-containing CIS cargo protein.

## References

1. Porello, I. & Cellesi, F. Intracellular delivery of therapeutic proteins. New advancements and future directions. Front. Bioeng. Biotechnol. 11, 1211798 (2023).

2. Shoari, A., Tooyserkani, R., Tahmasebi, M. & Löwik, D. W. P. M. Delivery of Various Cargos into Cancer Cells and Tissues via Cell-Penetrating Peptides: A Review of the Last Decade. Pharmaceutics 13, 1391 (2021).

3. Yuan, Y. et al. Intelligent Design of Lipid Nanoparticles for Enhanced Gene Therapeutics. Mol. Pharm. 22, 1142–1159 (2025).

4. Vora, S. et al. Rational design of a compact CRISPR-Cas9 activator for AAV-mediated delivery. bioRxiv 298620 (2018) doi:10.1101/298620.

5. Mangeot, P. E., Guiguettaz, L., Sohier, T. J. M. & Ricci, E. P. Delivery of the Cas9/sgRNA Ribonucleoprotein Complex in Immortalized and Primary Cells via Virus-like Particles (“Nanoblades”). J. Vis. Exp. (2021) doi:10.3791/62245-v.

6. Bai, F., Li, Z., Umezawa, A., Terada, N. & Jin, S. Bacterial type III secretion system as a protein delivery tool for a broad range of biomedical applications. Biotechnol. Adv. 36, 482–493 (2018).

7. Guzmán-Herrador, D. L., Fernández-Gómez, A. & Llosa, M. Recruitment of heterologous substrates by bacterial secretion systems for transkingdom translocation. Front. Cell. Infect. Microbiol. 13, 1146000 (2023).

8. Wettstadt, S. & Filloux, A. Manipulating the type VI secretion system spike to shuttle passenger proteins. PLoS ONE 15, e0228941 (2020).

9. Kreitz, J. et al. Programmable protein delivery with a bacterial contractile injection system. Nature 616, 357–364 (2023).

10. Kreitz, J. et al. Targeted delivery of diverse biomolecules with engineered bacterial nanosyringes. Nat. Biotechnol. 1–5 (2025) doi:10.1038/s41587-025-02774-x.

11. Legendre, M. G., Heredia, C. A., Colee, C. & Demirer, G. S. Extracellular contractile injection systems for high efficiency protein delivery to plants. bioRxiv 2025.08.22.671880 (2025) doi:10.1101/2025.08.22.671880.

12. Xu, J. et al. Identification and structure of an extracellular contractile injection system from the marine bacterium Algoriphagus machipongonensis. Nat Microbiol 7, 397–410 (2022).

13. Xu, J., Ericson, C. F., Toenshoff, E. R. & Pilhofer, M. Stepwise firing mechanism of an extracellular contractile injection system. bioRxiv 2025.10.08.681156 (2025) doi:10.1101/2025.10.08.681156.

14. Hyman, P. & Raaij, M. van. Bacteriophage T4 long tail fiber domains. Biophys. Rev. 10, 463–471 (2018).

15. Williams, S. R., Gebhart, D., Martin, D. W. & Scholl, D. Retargeting R-Type Pyocins To Generate Novel Bactericidal Protein Complexes. Appl Environ Microb 74, 3868–3876 (2008).

16. Hu, B., Margolin, W., Molineux, I. J. & Liu, J. Structural remodeling of bacteriophage T4 and host membranes during infection initiation. Proc National Acad Sci 112, E4919–E4928 (2015).

17. Weiss, G. L. et al. Structure of a thylakoid-anchored contractile injection system in multicellular cyanobacteria. Nat Microbiol 7, 386–396 (2022).

18. Abramson, J. et al. Accurate structure prediction of biomolecular interactions with AlphaFold 3. Nature 630, 493–500 (2024).

19. Garcia-Doval, C. & Raaij, M. J. van. Structure of the receptor-binding carboxy-terminal domain of bacteriophage T7 tail fibers. Proc. Natl. Acad. Sci. 109, 9390–9395 (2012).

20. Katoh, Y., Nozaki, S., Hartanto, D., Miyano, R. & Nakayama, K. Architectures of multisubunit complexes revealed by a visible immunoprecipitation assay using fluorescent fusion proteins. J. Cell Sci. 128, 2351–2362 (2015).

21. Adler, H. I., Fisher, W. D., Cohen, A. & Hardigree, A. A. MINIATURE escherichia coli CELLS DEFICIENT IN DNA*. Proc. Natl. Acad. Sci. 57, 321–326 (1967).

22. Farley, M. M., Hu, B., Margolin, W. & Liu, J. Minicells, Back in Fashion. J. Bacteriol. 198, 1186–1195 (2016).

23. Zhang, Y. & Bergelson, J. M. Adenovirus Receptors. J. Virol. 79, 12125–12131 (2005).

24. Jiang, F. et al. N-terminal signal peptides facilitate the engineering of PVC complex as a potent protein delivery system. Sci Adv 8, eabm2343 (2022).

25. Ericson, C. F. et al. A contractile injection system stimulates tubeworm metamorphosis by translocating a proteinaceous effector. Elife 8, e46845 (2019).

26. Vlisidou, I. et al. The Photorhabdus asymbiotica virulence cassettes deliver protein effectors directly into target eukaryotic cells. Elife 8, e46259 (2019).

27. Chen, L. et al. Genome-wide Identification and Characterization of a Superfamily of Bacterial Extracellular Contractile Injection Systems. Cell Reports 29, 511–521.e2 (2019).

28. Geller, A. M. et al. The extracellular contractile injection system is enriched in environmental microbes and associates with numerous toxins. Nat Commun 12, 3743 (2021).

29. Böck, D. et al. In situ architecture, function, and evolution of a contractile injection system. Science 357, 713–717 (2017).

30. Forster, A. et al. Coevolution of the ATPase ClpV, the TssB-TssC Sheath and the Accessory HsiE Protein Distinguishes Two Type VI Secretion Classes. (2014) doi:10.2210/pdb4uqw/pdb.

31. Douzi, B. et al. Structure and specificity of the Type VI secretion system ClpV-TssC interaction in enteroaggregative Escherichia coli. Sci. Rep. 6, 34405 (2016).

32. Kapitein, N. et al. Function of the AAA+ protein ClpV in type VI protein secretion. Mol Microbiol 87, 1013–1028 (2013).

33. Wendler, P., Ciniawsky, S., Kock, M. & Kube, S. Structure and function of the AAA+ nucleotide binding pocket. Biochim. Biophys. Acta (BBA) - Mol. Cell Res. 1823, 2–14 (2012).

34. Rybakova, D., Schramm, P., Mitra, A. K. & Hurst, M. R. H. Afp14 is implicated in the determination of Afp length. Mol Microbiol 96, 815–826 (2015).

35. Mahony, J. et al. Functional and structural dissection of the tape measure protein of lactococcal phage TP901-1. Sci. Rep. 6, 36667 (2016).

36. Wu, S., Liu, B. & Zhang, X. Identification of a tail assembly gene cluster from deep-sea thermophilic bacteriophage GVE2. Virus Genes 38, 507–514 (2009).

37. Xu, J., Hendrix, R. W. & Duda, R. L. Chaperone–Protein Interactions That Mediate Assembly of the Bacteriophage Lambda Tail to the Correct Length. J. Mol. Biol. 426, 1004–1018 (2014).

38. Cavrois, M., Noronha, C. de & Greene, W. C. A sensitive and specific enzyme-based assay detecting HIV-1 virion fusion in primary T lymphocytes. Nat. Biotechnol. 20, 1151–1154 (2002).

39. Jones, D. M. & Padilla-Parra, S. The β-Lactamase Assay: Harnessing a FRET Biosensor to Analyse Viral Fusion Mechanisms. Sensors 16, 950 (2016).

40. Zlokarnik, G. et al. Quantitation of Transcription and Clonal Selection of Single Living Cells with β-Lactamase as Reporter. Science 279, 84–88 (1998).

41. Cunliffe, T. G., Parker, A. L. & Jaramillo, A. Pseudotyping Bacteriophage P2 Tail Fibers to Extend the Host Range for Biomedical Applications. ACS Synth. Biol. 11, 3207–3215 (2022).

42. Dams, D., Brøndsted, L., Drulis-Kawa, Z. & Briers, Y. Engineering of receptor-binding proteins in bacteriophages and phage tail-like bacteriocins. Biochem. Soc. Trans. 47, 449–460 (2018).

43. Danov, A. et al. Identification of novel toxins associated with the extracellular contractile injection system using machine learning. Mol. Syst. Biol. 20, 859–879 (2024).

44. Puchades, C., Sandate, C. R. & Lander, G. C. The molecular principles governing the activity and functional diversity of AAA+ proteins. Nat. Rev. Mol. Cell Biol. 21, 43–58 (2020).

45. Olivares, A. O., Baker, T. A. & Sauer, R. T. Mechanistic insights into bacterial AAA+ proteases and protein-remodelling machines. Nat. Rev. Microbiol. 14, 33–44 (2016).

46. Katsura, I. & Hendrix, R. W. Length determination in bacteriophage lambda tails. Cell 39, 691–698 (1984).

47. Zhou, Z. H. et al. Atomic structures of a bacteriocin targeting Gram-positive bacteria. Res. Sq. rs.3.rs-4007122 (2024) doi:10.21203/rs.3.rs-4007122/v1.

48. Linares, R. et al. Structural basis of bacteriophage T5 infection trigger and E. coli cell wall perforation. Sci. Adv. 9, eade9674 (2023).

49. Demichev, V., Messner, C. B., Vernardis, S. I., Lilley, K. S. & Ralser, M. DIA-NN: Neural networks and interference correction enable deep proteome coverage in high throughput. Nat. methods 17, 41–44 (2020).

50. Wolski, W. E. et al. prolfqua: A Comprehensive R-Package for Proteomics Differential Expression Analysis. J. Proteome Res. 22, 1092–1104 (2023).

51. Huber, W., Heydebreck, A. von, Sültmann, H., Poustka, A. & Vingron, M. Variance stabilization applied to microarray data calibration and to the quantification of differential expression. Bioinformatics 18, S96–S104 (2002).

52. Feldmüller, M. et al. Stepwise assembly and release of Tc toxins from Yersinia entomophaga. Nat. Microbiol. 9, 405–420 (2024).

53. Punjani, A., Rubinstein, J. L., Fleet, D. J. & Brubaker, M. A. cryoSPARC: algorithms for rapid unsupervised cryo-EM structure determination. Nat. Methods 14, 290–296 (2017).

54. Casu, B., Sallmen, J. W., Schlimpert, S. & Pilhofer, M. Cytoplasmic contractile injection systems mediate cell death in Streptomyces. Nat. Microbiol. 8, 711–726 (2023).

55. Afanasyev, P. et al. A posteriori correction of camera characteristics from large image data sets. Sci. Rep. 5, 10317 (2015).

56. Kremer, J. R., Mastronarde, D. N. & McIntosh, J. R. Computer Visualization of Three-Dimensional Image Data Using IMOD. J. Struct. Biol. 116, 71–76 (1996).

57. Hunter, S. et al. InterPro: the integrative protein signature database. Nucleic Acids Res.37, D211–D215 (2009).

58. Eddy, S. R. Accelerated Profile HMM Searches. PLoS Comput. Biol. 7, e1002195 (2011).

59. Mistry, J. et al. Pfam: The protein families database in 2021. Nucleic Acids Res. 49, D412–D419 (2020).

60. Mészáros, B., Erdős, G. & Dosztányi, Z. IUPred2A: context-dependent prediction of protein disorder as a function of redox state and protein binding. Nucleic Acids Res. 46, W329–W337 (2018).

61. Kuriata, A., et al. CABS-flex 2.0: a web server for fast simulations of flexibility of protein structures. arXiv (2018) doi:10.48550/arxiv.1802.07568.

62. Kumar, S., Stecher, G., Li, M., Knyaz, C. & Tamura, K. MEGA X: Molecular Evolutionary Genetics Analysis across Computing Platforms. Mol. Biol. Evol. 35, 1547–1549 (2018).

